# An early, novel arginine methylation of KCa3.1 attenuates subsequent T cell exhaustion

**DOI:** 10.1101/2024.05.09.593421

**Authors:** Piyush Sharma, Ao Guo, Suresh Poudel, Emilio Boada-Romero, Katherine C. Verbist, Gustavo Palacios, Kalyan Immadisetty, Mark J. Chen, Dalia Haydar, Ashutosh Mishra, Junmin Peng, M. Madan Babu, Giedre Krenciute, Evan S. Glazer, Douglas R. Green

## Abstract

T cell receptor (TCR) engagement initiates the activation process, and this signaling event is regulated in multifaceted ways. Nutrient availability in the immediate niche is one such mode of regulation^1–3^. Here, we investigated how the availability of an essential amino acid methionine (Met) and TCR signaling might interplay in the earliest events of T cell activation to affect subsequent T cell fate and function. We found that limiting Met during only the initial 30 minutes of CD8^+^ T cell activation increased Ca^2+^ influx, Ca^2+^-mediated NFAT1 (*Nfatc2*) activation, NFAT1 promoter occupancy, and T cell exhaustion. We identified changes in the protein arginine methylome during the initial 30 min of TCR engagement and discovered a novel arginine methylation of a Ca^2+^-activated potassium transporter, KCa3.1, which regulates Ca^2+^-mediated NFAT1 signaling to ensure optimal activation. Ablation of arginine methylation in KCa3.1 led to increased NFAT1 activation, rendering T cells dysfunctional in murine tumour and infection models. Furthermore, acute Met supplementation at early stages reduced nuclear NFAT1 in tumour-infiltrating T cells and augmented their anti-tumour activity. Our findings identify a metabolic event occurring early after T cell activation that influences the subsequent fate of the cell.

## Methionine limitation promotes T cell exhaustion

During TCR engagement, nutrient metabolism ensures effective and optimal T cell activation and function^4^. To determine which amino acids were rapidly consumed upon T cell activation, we activated OT-I transgenic CD8^+^ T cells in complete medium and performed targeted mass spectrometry. We observed a significant decrease in intracellular Met and other amino acids as early as 10 min (Fig. 1A). Among the amino acids important for T cells, Met is the sole amino acid responsible for cellular methylome maintenance via the S-adenosyl Met (SAM) pathway, and alterations in the SAM pathway adversely affect T cell functions^5,6^. In complete media, no concomitant changes were observed in SAM or S-adenosyl homocysteine (SAH) (Fig. 1B, upper panel). T cells consumed their intracellular Met pool only in Met-deprived medium, correlating with decreases in SAM concentration and corresponding increases in SAH (Fig.1B, bottom panel). These observations indicate that extracellular Met availability is required to maintain the SAM cycle during T cell activation.

**Figure 1:**
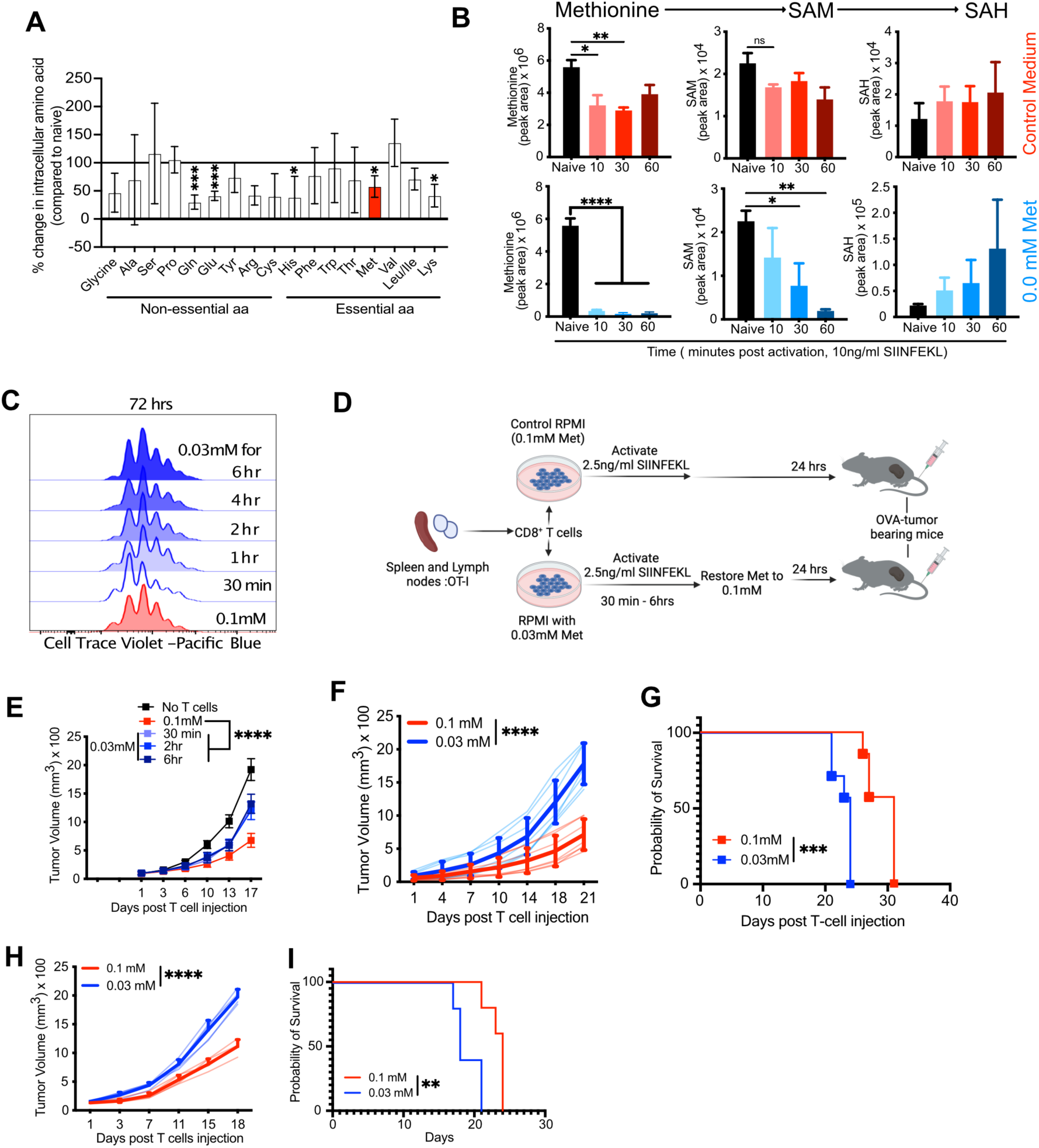
TCR-mediated, rapid methionine consumption governs T cell effector function. **a-b,** Quantification of intracellular amino acids at 10 min (a) and SAM and SAH up to 60 min (b) in OT-I T cells activated with 10 ng/ml SIINFEKL (n=3). **c,** T cell proliferation via cell-trace violet staining of OT-I CD8^+^ T cells activated in either 0.1 mM Met or 0.03 mM Met for the indicated times before restoration to 0.1 mM Met in 0.03 mM Met conditions and then analyzed 72 hrs post activation (n=3). Representative of 3 samples per group. **d,** Schematic design for OT-I T cell activation initially in 0.1 or 0.03 mM Met, followed by restoration of Met to 0.1 mM for 24 hrs before injection into B16-OVA tumour-bearing mice. **e,** Tumour growth of B16-OVA in *Rag1*^-/-^ treated with OT-I CD8^+^ T cells activated as described in (d) for 30 min-6hrs (n=5). **f-g,** Tumour growth (f) and survival (g) of B16-OVA tumours in *Rag1*^-/-^ mice after transfer of 24 hr-activated OT-I T cells with first 30 min of stimulation being in 0.1 or 0.03 mM Met with 2.5ng/ml SIINFEKL before restoration to 0.1 mM Met (n=5). **h-i,** Tumour growth (h) and survival (i) of B16-OVA tumours treated with activated GP33^+^-memory T cells as described in extended data fig. 1d (n=5). Data are mean±s.e.m. Unpaired two-tailed Student’s t-test (a, b), two-way ANOVA (e, f, h), Mantel-Cox log rank test (g, i) and paired two-tailed Student’s t-test (G-L) *P < 0.05, **P < 0.01, ***P < 0.001, ****P < 0.0001.

Mice infected with chronic Lymphocytic choriomeningitis virus (LCMV) have ∼65% less Met than in serum as early as day 2 post infection^7^ (Extended Data Fig. 1A), suggesting that T cells can experience low Met conditions during activation. To determine whether Met limitation during TCR engagement affects T cell proliferation, we lowered the available amount of Met to 0.03 mM in the culture medium. Cell-trace violet (CTV) labelled OT-I CD8^+^ T cells were activated in media with control (0.1 mM) or 0.03 mM Met for 30 min to 6 hr, after which Met was restored to 0.1 mM, and we observed no difference in the proliferation of T cells activated in reduced Met for different times (Fig. 1C). When then assessed their activity *in vivo* upon transferring these activated T cells into mice bearing B16/MC38 tumours expressing ovalbumin (B16-OVA/ MC38-OVA) 24 h after activation (Fig. 1D), we observed defective tumour control in T cells activated in 0.03 mM Met compared to 0.1 mM Met, across all time points, with the earliest being 30 min (*Rag1*^-/-^ mice: Fig. 1E-G-B16-OVA; B6 WT mice: Extended Data Fig. 1B, C-MC38-OVA). To further assess this effect, we generated LCMV GP33-specific memory T cells and activated them for 24 hrs with GP33 peptide in either 0.1 mM Met or 0.03 mM Met for 30 min, after which Met was restored to 0.1 mM and then transferred them into B16-GP33 (B16 expressing GP33 peptide) tumour-bearing *Rag1^-/-^* mice (Extended Data Fig. 1D). We observed defective tumour control leading to poor survival in mice receiving T cells activated in 0.03 mM Met compared to 0.1 mM Met (Fig. 1H and I).

To examine the effects of activation in reduced Met on the T cell epigenome, we performed ATAC-Seq at 24 hrs post activation (Extended Data Fig. 1E) and observed increased chromatin accessibility in T cells activated in 0.03 mM Met during the initial 30 min (Extended Data Fig. 2A and B). Overrepresentation analysis showed enrichment of gene sets associated with an exhaustion-like phenotype (Extended Data Fig. 2C). Further HOMER motif analysis showed that transcription factor (TF) motifs known to be associated with exhausted T cells^8,9^ were highly accessible in 0.03 mM-activated T cells (Extended Data Fig. 2D).

We activated OT-I CD8^+^ T cells T cells in either control or 0.03 mM Met for the initial 30 min followed by 24 hours culture in 0.1 mM Met and injected them into B16-OVA bearing *Rag1^-/-^* mice. TILs were collected at day 9 post T cell injection. Again, animals receiving T cells activated initially in 0.03 mM Met showed defective tumour control (Extended Data Fig. 2E and F). The CD8^+^ TILs displayed reduced CD62L^+^, CD62L^high^ CD44^high^ central memory cells (Tcm) with increased TOX expression and reduced IFNγ production compared to TILs from T cells activated in Met replete media (Extended Data Fig. 2G).

T cell dysfunction is known to contribute to poor prognoses in cancer^10^. This dysfunctional state is at least in part driven by the transcription factor (TF) TOX^11^ and is accompanied by markers such as increased surface expression of PD1 and Tim-3 and reduced effector cytokine production^12,13^. To further examine the effect of activation in limiting Met, we activated CD45.1^+^ or CD45.2^+^ OT-I CD8^+^ T cells in 0.1 (control) or 0.03 mM Met for 30 min, restored Met to 0.1 mM for 24 hrs, and mixed them at a ratio of 1:1 before transferring them into B16-OVA tumour-bearing *Rag1*^-/-^ mice (Fig. 2A). Even though CD8^+^ TIL initially activated in 0.03 mM outnumbered the 0.1 mM-activated CD8^+^ TIL at day 12 (Fig. 2B and C), these cells exhibited significant increases in exhaustion-associated expression of PD1 and Tim-3, reduced effector cytokines (Fig. 2D,E and Extended Data Fig. 2H), increased expression of exhaustion associated TF, TOX (Fig. 2F and G), and reduced expression of the stemness-associated TF, TCF-1 (Extended Data Fig. 2I). Altogether, these results suggest that reduced Met during early TCR signaling promotes acquisition of epigenomic changes associated with exhaustion and drives T cells towards a dysfunctional, exhausted state.

**Figure 2:**
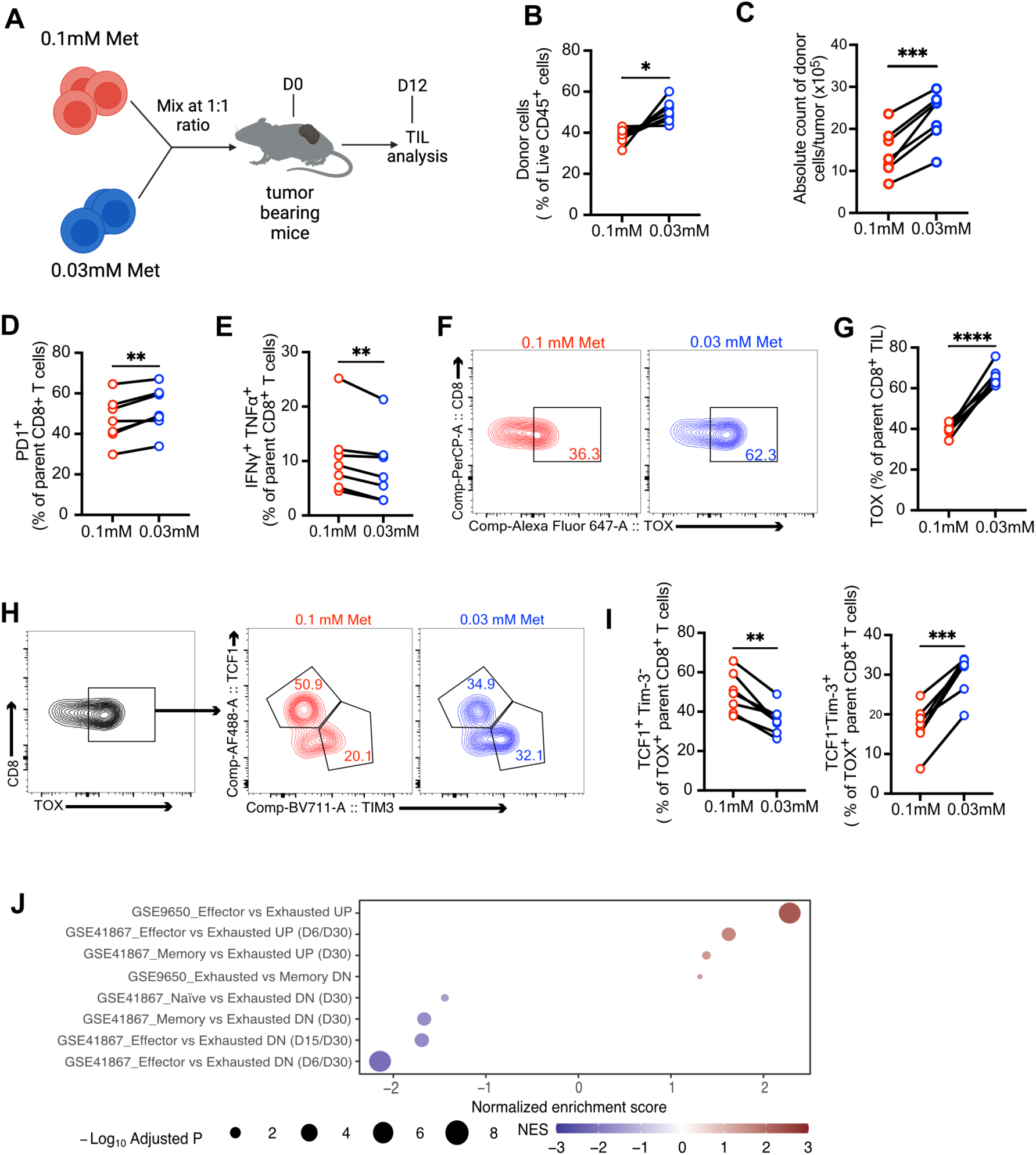
Reduced methionine availability during TCR signaling promotes T cell exhaustion. **a,** Schematic of experimental design. OT-I CD8^+^ T cells with different congenic markers were initially activated in 0.1 mM or 0.03 mM Met for 30 min with replenishment of Met in 0.03 mM to 0.1 mM Met for 24 hrs, transferred into B16-OVA tumour-bearing *Rag1*^-/-^ mouse at a 1:1 ratio, and analyzed day 12 post T cell transfer. **b-c,** Frequencies (b) and absolute number (c) of transferred T cells isolated from B16-OVA tumours at D12 post transfer (n=7). **d-e,** Frequency of PD1^+^ and IFNγ^+^TNFα^+^ OT-I CD8^+^ TIL, initially activated in 0.1 mM and 0.03 mM Met and assessed at D12 post injection (as in a) (n=7). **f-g,** Contour plot (f) and frequency of TOX^+^ cells (g) and of OT-I CD8^+^ TIL as in d (n=7). **h-i,** Representative contour plot (h) and quantification (i) of TCF1 and Tim-3 expressing cells from TOX^+^ CD8^+^ TIL from OT-I CD8^+^ T cells isolated at D12 post transfer as in (a) (n=7). **j,** T cell exhaustion gene sets in GSEA analysis of RNA-Seq of OT-I CD8^+^ TIL on d9 post T cell injection as in Fig. 1f (n=4). Data are mean±s.e.m. paired two-tailed Student’s t-test. *P < 0.05, **P < 0.01, ***P < 0.001.

Exhausted T cells have been further defined as precursors of exhaustion (TEXprog) and TEXprogenitor-like cells, both of which produce a replicative burst and effector function upon checkpoint blockade ^14^, and terminally exhausted cells (TEXterm), which fail to mount an effective effector response ^15,16^. Although TOX is associated with maintenance of exhausted T cells, it is also expressed in TEXprog ^17^. We therefore further dissected the TOX^+^ CD8^+^ T cells from the experiment in Fig. 2F (Fig. 2H) and found decreased TEXprog (TCF1^+^Tim-3^-^) and increased TEXterm (TCF1^-^Tim-3^+^) among T cells initially activated in 0.03 mM Met, suggesting a decreased pool of stem-like TEXprog cells (Fig. 2I). Furthermore, evaluation of total CD8^+^ TILs from mice receiving T cells initially activated in 0.03 mM Met for 30 minutes revealed a lower frequency of TCF1^+^Tim-3^-^ TEXprog cells and CX3CR1^+^Ly108^+^ TEXprogenitor-like cells (Extended Data Fig. 2J) and a corresponding increase in the frequency of TEXterm cells defined as TCF1^-^Tim-3^+^ (Extended Data Fig. 2K). We also observed a decrease in memory-associated markers such as CD62L^+^, CD62L^+^CD44^+^ (Extended Data Fig. 2L) and CD127^+^CD27^+^ (Extended Data Fig. 2M and N) and an increase in KLRG1^+^CD127^-^ terminal effector-like T cells (Extended Data Fig. 2O and P) when initially activated in 0.03 mM Met.

To examine the transcriptomic state, we performed global RNA-seq of day 9 TIL from T cells originally activated for 30 min in 0.1 mM or 0.03 mM Met and then cultured in 0.1 mM Met for 24hrs prior to transfer into B16-OVA tumour-bearing *Rag1-/-* mice. We found several transcriptomic changes, the majority of which represented increased expression of differentially enriched (DE) genes in T cells initially activated in 0.03 mM Met (Extended Data Fig. 3A and B).

Over-representation analysis of the DE genes in T cells activated in 0.1 mM Met revealed enrichment of GOBP pathways which are associated with T cell effector functions along with increases in transcription factors associated with a stem-like T cell state and memory generation ^18–21^ (Extended Data Fig. 3C). Cells initially activated in 0.03 mM Met showed enrichment of genes which are associated with terminal differentiation or dysfunction of T cells and transcription factors associated with hyperactivation and development and maintenance of T cell exhaustion ^8,22–24^ (Extended Data Fig. 3D). GSEA analysis also showed an enriched exhaustion-associated gene set ^25,26^ in CD8^+^ TIL initially activated in 0.03 mM Met (Fig. 2J) and enrichment of hallmark gene sets associated with T cell effector function in the control (0.1 mM Met) T cells (Extended Data Fig. 3E). These results suggest that Met availability during early TCR engagement influences T cell differentiation, with initial, reduced Met availability promoting hyperactivation and driving T cells towards exhaustion.

## Extracellular Met metabolism regulates Ca^2+^ flux and NFAT1 activation

Among the earliest events in T cell activation is the influx of Ca^2+^ ^27,28^, which is accompanied by the activation of calcineurin, which dephosphorylates Nuclear Factor of Activated T cells-1 (NFAT1), resulting in its nuclear translocation and activation^29,30^. Because low Met levels for only the first 30 min of activation influenced the subsequent T cell phenotype, we investigated changes in Ca^2+^ flux using Fluo-8. We observed increased Ca^2+^ influx upon stimulation of T cells with anti-CD3/CD28 antibody in 0.03 mM compared to 0.1 mM Met (Fig. 3A), which was reduced by the CRAC channel inhibitor YM-58483 (Extended Data Fig. 4A). NFAT1 cooperates with the AP1 complex to ensure optimal T cell activation and differentiation, but high levels of NFAT1 activity have been associated with T cell dysfunction and exhaustion ^29,31^. Confocal imaging revealed an increase in activated NFAT1 (nuclear localization of NFAT1) in T cells activated for 30 min with anti-CD3/28 beads in 0.03 mM Met compared to 0.1 mM Met (Fig. 3B, Extended Data Fig. 4B), quantified as either a ratio of nuclear NFAT1 to total cell NFAT (Fig. 3C and Extended Data Fig. 4C(0.0 mM Met)) or as nuclear NFAT1 fluorescence intensity (Extended Data Fig. 4D), which were abolished upon treatment with either the calcineurin inhibitor Cyclosporin A (CsA) (Fig. 3B and C) or CRAC channel inhibitor YM-58483 (Extended Data Fig. 4E). Examination of NFAT2 at 30 min of activation revealed no difference in its activation in T cells activated in either 0.1 mM or 0.03 mM Met (Extended Data Fig. 4F). Next, we activated CD8^+^ T cells in complete medium for 24hrs and rested them for 48hrs before secondary activation in 0.03 mM or 0.1 mM Met and observed increased nuclear NFAT1 levels in 0.03 mM Met 30 min after activation (Extended Data Fig. 4G), suggesting that the effect of limiting Met on NFAT1 activation is independent of previous activation events.

**Figure 3:**
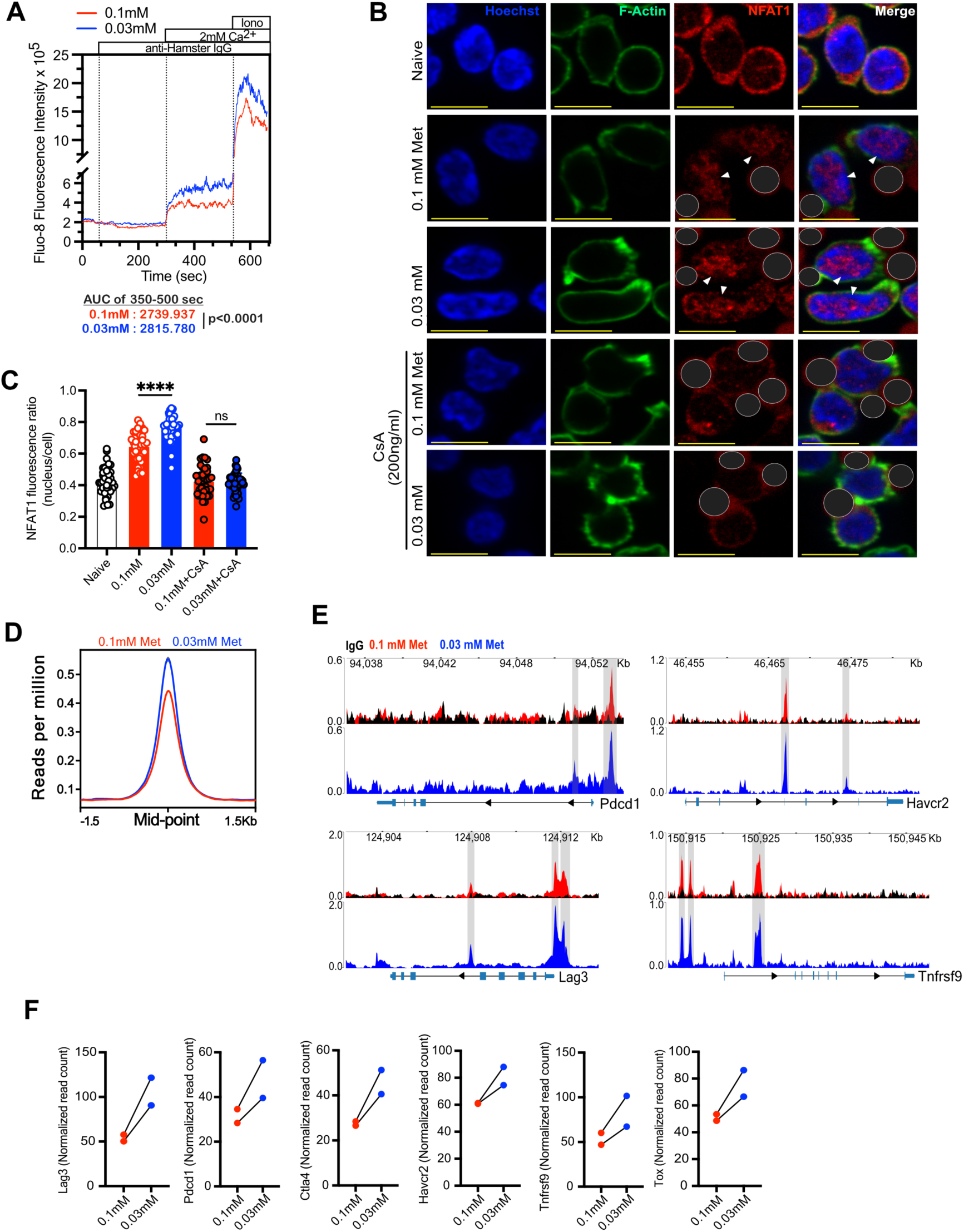
Extracellular methionine availability regulates Ca^2+^-mediated NFAT1 activity. **a,** Fluo-8 AM analysis of Ca^2+^ flux in CD8^+^ T cells activated with anti-CD3 and anti-CD28 by anti-hamster IgG crosslinking in either 0.1 mM or 0.03 mM Met containing Ca^2+^ Ringer solution with addition of 2 mM Ca^2+^ to measure Ca^2+^ influx. The area under the curve is measured from 350-500 sec. Ionomycin then added as a positive control for maximal flux. Representative of 2 experiments. **b,** Representative images of T cells, cultured with or without anti-CD3/28 dynabeads (dark grey masked), in 0.1 mM and 0.03 mM Met, with or without cyclosporin-A treatment for 30 min, stained with NFAT1 (red, marked), Hoechst (blue) and Phalloidin (green). **c,** Quantification of NFAT1 intensity as nuclear to total cell ratio (c) (See Fig. S4A for unmasked figure). Scale bar=10μm. (Each circle represents one cell, n=40 cells). **d,** Histogram of NFAT1 binding signals (read count per million reads normalized to background) from NFAT1 CUT&RUN on T cells initially activated in 0.1 or 0.03 mM Met for 30 min and assessed 24 hrs post activation. **e-f,** NFAT1 CUT&RUN peaks at the known target genes (e) and quantification of known NFAT1-binding regions of *Lag3, Pdcd1, Havcr2, Ctla4* and *Tnfrsf9 and Tox* (f, normalized read count (see differential binding analysis)) in T cells initially activated in 0.1 or 0.03 mM Met for 30 min and assessed 24 hrs post activation (n=2). Data are mean±s.e.m. Unpaired two-tailed Student’s t-test (C, D). *P < 0.05, **P < 0.01, ***P < 0.001, ****P < 0.0001.

NFAT1 is known to regulate several exhaustion-associated genes such as *Lag3, Pdcd1, Havcr2, Ctla4, Tnfrsf9* ^31^. We performed NFAT1 CUT&RUN on T cells activated in 0.1 mM or 0.03 mM Met for 30 min, followed by 0.1 mM Met for 24hrs and observed increased NFAT1 binding in the low Met group (Fig. 3D). We observed that Met limitation for the initial 30 min resulted in increased NFAT1 binding to *Pdcd1, Havcr2, Lag3, and Tnfrsf9* (Fig. 3E) as well as *Ctla4 and Tox* gene loci (Fig. 3F). This further correlated with increased transcription of most of these genes at 24 hrs (Extended Data Fig. 4H). Altogether, our results suggest that reduced Met availability during the initial stages of TCR activation directs a trajectory of T cells towards hyperactivation and eventual exhaustion, and the maintenance of extracellular Met levels in the face of rapid Met consumption upon TCR engagement dampens Ca^2+^-mediated NFAT1 activity and T cell fate.

## KCa3.1 methylation regulates NFAT1

Methionine metabolism is the sole pathway for epigenetic and proteomic methylome maintenance, which is known to affect T cell function ^32^. When converted to SAH, SAM acts as the methyl donor; thus, the ratio of SAM to SAH is known as the cellular methylation potential ^33^, an index of a cell’s capacity to maintain its methylome. We calculated the SAM/SAH ratio over time after TCR engagement in the absence of extracellular Met and found this ratio to be significantly reduced as early as 10 min post-activation (Extended Data Fig. 5A), suggesting alterations in the methylome. Met metabolism is essential for DNA methylation ^34^; however, we found no significant differences in global 5-methylcytosine levels at either 30 min or 24 hrs in T cells activated as above (Extended Data Fig. 5B). We similarly assessed H3me3K27 and H3me2K4 by CUT&RUN at 24 hrs post activation, but again, no global changes were observed (Extended Data Fig. 5C and D). Thus, Met restriction for only the first 30 min of T cell activation does not appear to directly affect global DNA or histone methylation.

Therefore, we postulated that Met restriction influences early methylation events independently of direct effects on DNA and histone methylation. Post-translational protein arginine methylation mediated by protein arginine methyltransferases regulates protein function ^35,36^. We found that arginine methylation in T cells activated in 0.0 mM or 0.03 mM Met was significantly reduced 30 min post-activation when compared to T cells activated in 0.1 mM Met (Fig. 4A and B, Extended Data Fig. 5E). To identify specific proteins differentially methylated upon TCR engagement in limited Met, CD8^+^ T cells activated by anti-CD3 antibody with IgG cross-linking for 30 min were processed for methyl-arginine immunoprecipitation and quantification using multiplexed tandem mass tag-mass spectrometry (TMT-MS). We identified common proteins which were differentially methylated in 0.0 mM and 0.03 mM Met activation conditions compared to 0.1 mM Met (Fig. 4C) and found the top candidate, KCa3.1 to be di-methylated at Arginine-350 (R350) in 0.1 mM Met, which was reduced in 0.0 mM and 0.03 mM Met (Fig. 4D, Supplementary Table. 1). Furthermore, protein-BLAST analysis showed that KCa3.1 R350 is conserved across species, including humans, which suggested that R350 may be important for KCa3.1 regulation (Extended Data Fig. 5F).

**Figure 4.**
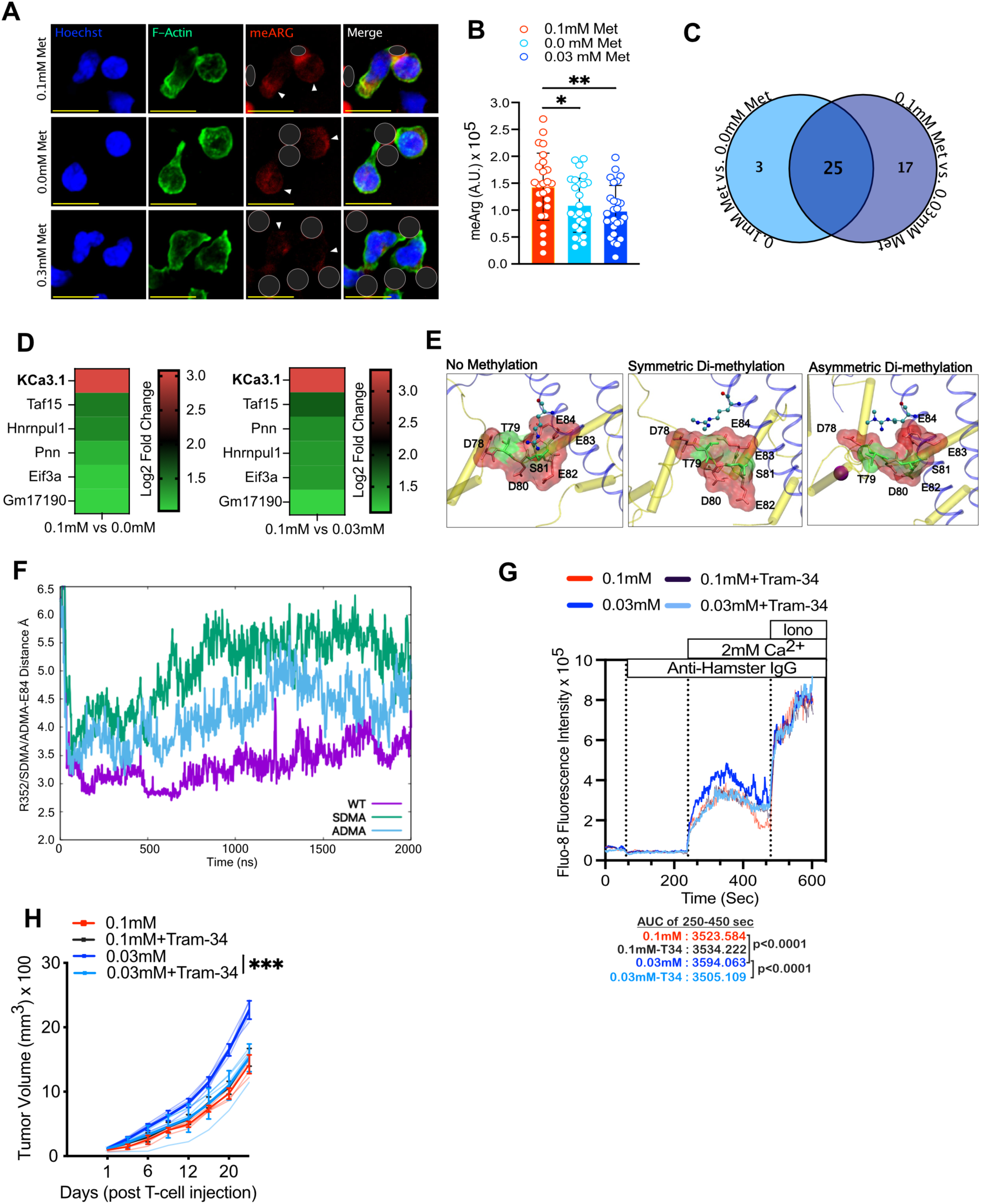
Methionine mediated KCa3.1 methylation upon TCR activation regulates T cell effector function. **a-b,** Representative images (a) and quantification (b) of methyl-arginine in CD8^+^ T cells activated with anti-CD3/28 dynabeads (circled in red channel images) in 0.1, 0.0, and 0.03 mM Met for 30 min, stained with pan-meArg (red, with arrows), Hoechst (blue), and Phalloidin (green) (scale bar=10 μm) (each circle represents one cell. n= 25 cells). **c,** Venn diagram of enriched methyl-arginine proteins at 30 min post activation of T cells in 0.1 mM vs 0.0 mM or 0.1 mM vs 0.03 mM Met, as identified by TMT-MS. **d,** Heat-map (Log_2_ fold change) of TMT-MS identified proteins (p<0.05) with enriched methyl-arginine in T cells activated in 0.1 mM vs 0.0mM or 0.03 mM Met by IgG crosslinking with anti-CD3 for 30 min (n=3). **e,** Predicted interaction of KCa3.1 monomer (blue cartoon) with calmodulin (yellow cartoon) as un-methylated (left), SDMA (center) and ADMA (right). The Calmodulin-binding pocket is represented as surface rendering with interacting amino acids as sticks (negative charges in red, polar residues in green) and R352 as ball and sticks; introduction of a hydrophobic methyl group (middle and bottom) weakens the interactions between calmodulin and the KCa3.1 monomer. **f,** MD simulation analysis: Analysis of variations of distance of salt bridge between calmodulin E84 and demethylated R352 WT (purple), SDMA R352 (green) and ADMA R352 (blue) over the course of simulation time (averaged across all four monomers, three MD trials). **g,** Fluo-8 AM analysis of Ca^2+^ flux in CD8^+^ T cells activated with anti-CD3 and anti-CD28 by anti-hamster IgG crosslinking in Ca^2+^-free Ringer solution supplemented with either 0.1 mM or 0.03 mM Met and treated with either with DMSO or 1μM TRAM-34. Area under the curve measured from 250-450 sec (representative of 2 experiments). **h,** Tumour growth of B16-OVA tumours in *Rag1*^-/-^ mouse after transfer of OT-I CD8^+^ T cells activated in 0.1 mM or 0.03 mM Met with either DMSO or 1 μM TRAM-34 for 30 min and cultured for 24 hrs (n=5). Data are mean±s.e.m. Unpaired two-tailed Student’s t-test (b), 2-way ANOVA (h), two-sample T-test (g) as described in the methods. *P < 0.05, **P < 0.01, ***P < 0.001.

KCa3.1 is a calcium-activated potassium transporter encoded by *Kcnn4* and is activated by calmodulin binding upon TCR-mediated Ca^2+^ flux^37^. During T cell activation, the plasma membrane is hyperpolarized to ensure sufficient Ca^2+^ flux for signaling^38^, and this is maintained by K^+^ efflux through potassium transporters such as KCa3.1^39,40^. Based on our earlier observation of increased Ca^2+^ signaling under low Met conditions, we hypothesized that R350 methylation of KCa3.1 regulates its activity, and loss of this methyl group may result in hyperactivation, leading to increased Ca^2+^ influx and NFAT1 activation. To understand how arginine methylation may mechanistically modulate KCa3.1 activity, we performed MD simulations with the solved crystal structure of human KCa3.1(hKCa3.1) bound to calmodulin (CaM) at R352 (corresponding to R350 in murine KCa3.1) (Extended Data Fig. 5G). The calmodulin-binding pocket encompasses D78-T79-D80-S81-E82-E83-E84, which is predominantly negatively charged/polar, where E84 forms a salt bridge with R352 of hKCa3.1. Our results showed that both symmetrical di-methylation (SDMA) and asymmetrical di-methylation (ADMA) of R352 introduced a hydrophobic methyl group into this negatively charged/polar CaM binding pocket, thereby weakening the hKCa3.1 and CaM interaction (Fig. 4E). Furthermore, MD simulations showed that the salt bridge interaction of R352 with E84 of the CaM binding pocket diminished over time, especially in the case of SDMA where both guanidino nitrogens are methylated, compared to ADMA in which only one nitrogen is methylated (Fig. 4F, Extended Data Fig. 5H), suggesting that disruption of arginine methylation promoted stabilization of calmodulin binding, leading to increased activation of the channel and Ca^2+^ flux.

To validate our *in-silico* simulation results, we used the chemical inhibitor TRAM-34 to reduce KCa3.1 activity ^41,42^ during T cell activation and observed reduced Ca^2+^ flux upon TRAM-34 treatment only in 0.03 mM Met and not in 0.1 mM Met (Fig. 4G) at the concentration of TRAM-34 we employed. Measurement of nuclear NFAT1 levels revealed that the treatment normalized nuclear NFAT1 levels in T cells activated in 0.03 mM Met in a dose-dependent manner and had no effect on T cells activated in 0.1 mM Met (Extended Data Fig. 5I). We then activated OT-I T cells in 0.1 mM or 0.03 mM Met with either DMSO or TRAM-34 for 30 min, washed and then cultured the cells for an additional 24 hrs in 0.1 mM Met before transferring into B16-OVA tumour-bearing *Rag1*^-/-^ mice (Extended Data Fig. 5J). Chemical inhibition of KCa3.1 for 30 min significantly improved anti-tumour activity in T cells activated for 30 min in 0.03 mM Met (Fig. 4H), suggesting that increased KCa3.1 activity was responsible for the eventual decreased T cell function under this activation condition.

## Methylation-deficient KCa3.1 increases Ca^2+^-mediated NFAT1 activation

To specifically interrogate the role of R350 methylation of KCa3.1 in T cell activation, we cloned KCa3.1 (Kca3.1^WT^) and a mutant with an alanine substitution (KCa.3.1^R350A^) into retroviral vectors and expressed them in primary murine CD8^+^ T cells in which endogenous KCa3.1 was deleted using CRISPR/Cas9 (Extended Data Fig. 6A, B and C), and expression of the two was similar (Extended Data Fig. 6D). Fluo-8 Ca^2+^ flux analysis showed significantly increased flux upon activation in KCa3.1^R350A^ T cells (Fig. 5A). Due to the expression of GFP by transduced T cells, which might interfere with the Fluo-8 signal, we used another Ca^2+^-sensitive dye, ICR-1AM, and observed similar results (Extended Data Fig. 6E). Increased Ca^2+^ flux correlated with an increase in nuclear NFAT1 30 min post activation (Fig. 5B, C and Extended Data Fig. 6F). This heightened nuclear NFAT1 in cells expressing KCa3.1^R350A^ was visible under resting conditions and increased upon activation in a time-dependent manner (Extended Data Fig. 6G).

**Figure 5.**
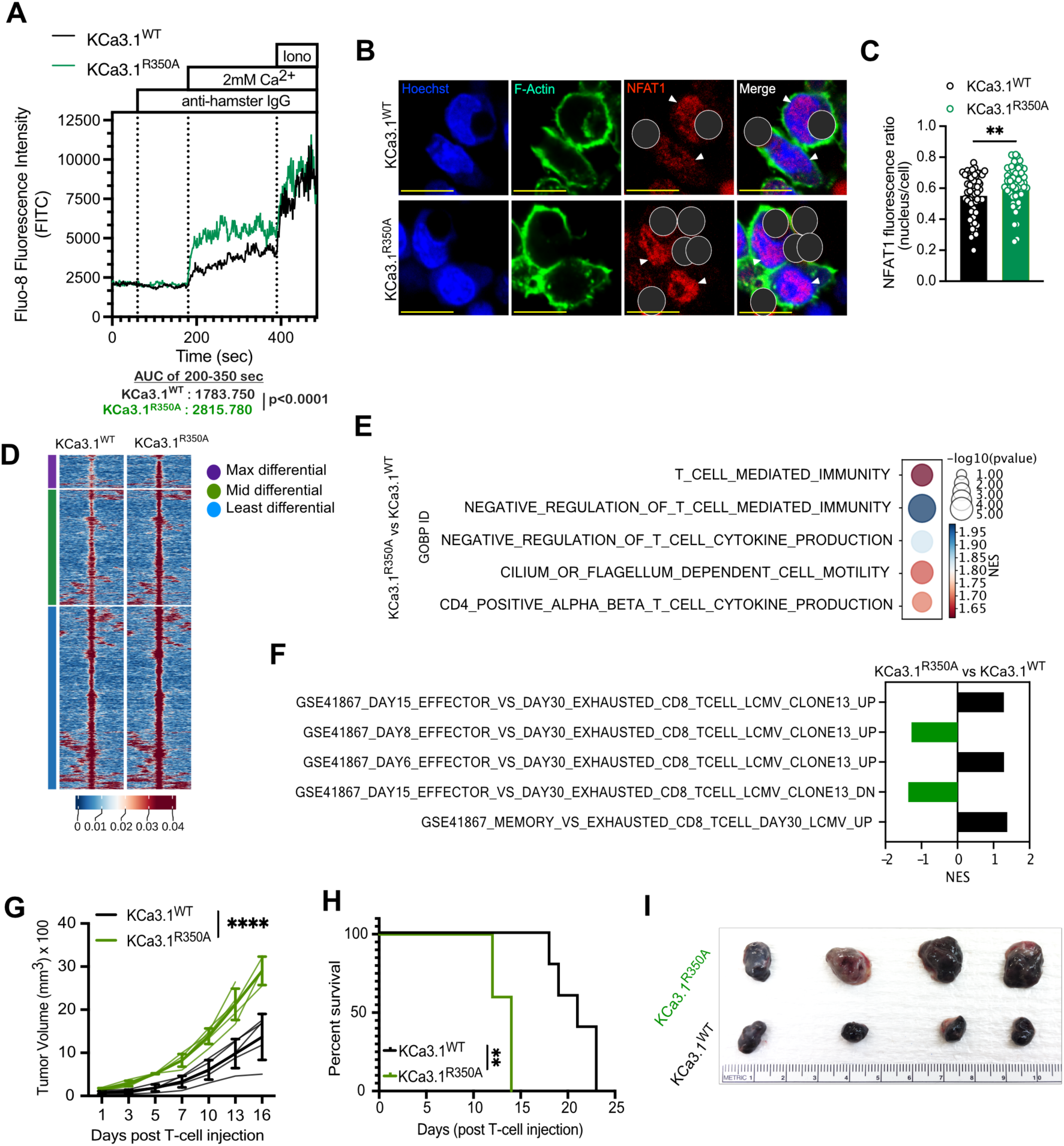
Ablation of KCa3.1 R350 methylation increased Ca^2+^-mediated NFAT1 activity, promoting T cell dysfunction. **a,** Fluo-8 analysis of Ca^2+^ flux in KCa3.1^WT^ and KCa3.1^R350A^ T cells activated with anti-CD3 and anti-CD28 by anti-hamster IgG crosslinking in Ca^2+^-free Ringer solution. Area under the curve calculated from 200-350 sec. **b-c,** Representative image (c) and quantification of NFAT1intensity as nuclear to total cell ratio (c) in KCa3.1^WT^ and KCa3.1^R350A^ OT-I T cells activated for 30 min with anti-CD3/28 dynabeads (dark grey masked) (See Extended Data Fig. 6F for unmasked figure), scale bar=10 μm (each circle represents one cell, n=40 cells). **d,** Chromatin accessibility heatmap of T cells expressing activated KCa3.1^WT^ or KCa3.1^R350A^ with each row representing peaks (p<0.05 and log2 fold-change >1.5) displayed over the span of 2kb window with peak as center (grouped from least to max differential region), analyzed 24 hrs post activation with 2.5ng/ml SIINFEKL (n=3). **e,** GOBP terms associated with negative regulation of T cells enriched in OT-I T cells expressing KCa3.1^R350A^ vs KCa3.1^WT^ activated as in (b) (n=3). **f,** T cell exhaustion gene sets in GSEA analysis of differential accessible promoter reads from ATAC-Seq of OT-I T cells expressing KCa3.1^WT^ or KCa3.1^R350A^, activated with 2.5 ng/ml SIINFEKL for 24 hrs (n=3). **g-h,** Tumour growth (g) and survival (h) of B16-OVA tumour-bearing RAG^-/-^ mice after transfer of KCa3.1^WT^ and KCa3.1^R350A^ OT-I T cells (n=5). **i,** B16-OVA tumours harvested at D12 post transfer of KCa3.1^WT^ and KCa3.1^R350A^ OT-I T cells (n=4). Data are mean±s.e.m. Two-sample T-test (a) as described in the methods, unpaired two-tailed Student’s t-test (c), 2-way ANOVA (g), and Mantel-Cox log rank test (h). **P < 0.01, ****P < 0.0001.

To investigate the epigenomic state of WT or mutant KCa3.1 T cells, we analyzed global chromatin accessibility via ATAC-Seq after 24 hrs of *in vitro* activation. We found that the activated KCa3.1^R350A^ T cells showed distinct chromatin accessibility which was increased compared to KCa3.1^WT^ T cells (Fig. 5D, Extended Data Fig. 7A). Furthermore, GSEA analysis revealed enrichment of GOBP terms associated with negative regulators of T cell-mediated immunity and gene sets associated with T cell exhaustion ^25,26^ in T cells expressing mutant KCa3.1^R350A^ (Fig. 5E and F), suggesting that ablation of R350 methylation drives T cells towards exhaustion early after activation. Upon comparing these ATAC-Seq results with ATAC-Seq of T cells activated in 0.03 mM vs. 0.1 mM Met (Extended Data Fig. 2B), we found several common differentially accessible regions (DAR) (Extended Data Fig. 7B) with enrichment of gene sets associated with T cell exhaustion (Extended Data Fig. 7C). HOMER motif analysis of WT or mutant T cells and common DARs was similarly enriched (Extended Data Fig. 2D, 7D and E). Despite these results, KCa.3.1^R350A^ did not fully phenocopy the chromatin effects of activation in 0.03 mM Met, suggesting that other differentially methylated proteins in addition to KCa.3.1 may also impact chromatin accessibility following activation in the latter condition.

To investigate whether the observed exhaustion-specific chromatin state led to defective functionality of mutant KCa3.1 T cells, we transferred T cells expressing KCa.3.1^WT^ or KCa.3.1^R350A^ into B16-OVA tumour-bearing Rag1^-/-^ mice. We found that T cells expressing KCa.3.1^R350A^ displayed impaired tumour control and survival (Fig. 5G, H and I), suggesting that the absence of KCa3.1 methylation drives T cells towards an exhaustion trajectory, leading to defective effector function and tumour control.

## Ablation of KCa3.1 methylation promotes T cell exhaustion

To determine whether ablation of R350 methylation drives T cell exhaustion in tumours, we transferred T cells expressing KCa.3.1^WT^ or KCa.3.1^R350A^ into B16-OVA tumour-bearing Rag1^-/-^ mice. Tumours were isolated and measured at day 12 post-T cell injection, and consistently, much larger tumours were extracted from animals treated with T cells expressing KCa.3.1^R350A^ (Extended Data Fig. 8A). KCa.3.1^R350A^-expressing TIL showed high surface expression of the exhaustion markers PD1, Lag3, and Tim-3 (Extended Data Fig. 8B) that correlated with increased TOX expression (Extended Data Fig. 8C). Among these cells we also found fewer TEXprog (CD69^lo^Ly108^hi^)^43^ and IFNγ- and TNFα-producing T cells (Extended Data Fig. 8D and E). Next, we performed a similar experiment in a competitive setting, wherein CD45.1^+^ KCa.3.1^WT^ and CD45.1^+^CD45.2^+^ Kca.3.1^R350A^-expressing OT-I CD8^+^ T cells were mixed at a 1:1 ratio and transferred into B16-OVA tumour-bearing *Rag1^-/-^* mice (Fig. 6A). TIL analysis at D12 showed increased PD1^+^Tim-3^+^, Eomes^+^PD1^+^ (Fig. 6B, Extended Data Fig. 8F) and TOX expression in Kca.3.1^R350A^-expressing T cells, which correlated with reduced expression of the effector cytokines IFNγ and TNFα (Fig. 6C and D). Further analysis revealed a decreased frequency of TCF1^+^Tim-3^-^ TEXprog and increased TCF1^-^Tim-3^+^ TEXterm populations (Extended Data Fig. 8G and Fig. 6E). We also observed a decrease in CD62L^+^ and CD127^+^CD27^+^ memory T cells (Fig. 6F) with an increase in the KLRG1^+^CD127^-^ terminal effector-like phenotype in KCa.3.1^R350A^-expressing T cells, as compared to T cells expressing WT KCa3.1 (Extended Data Fig. 8H and I).

**Figure 6.**
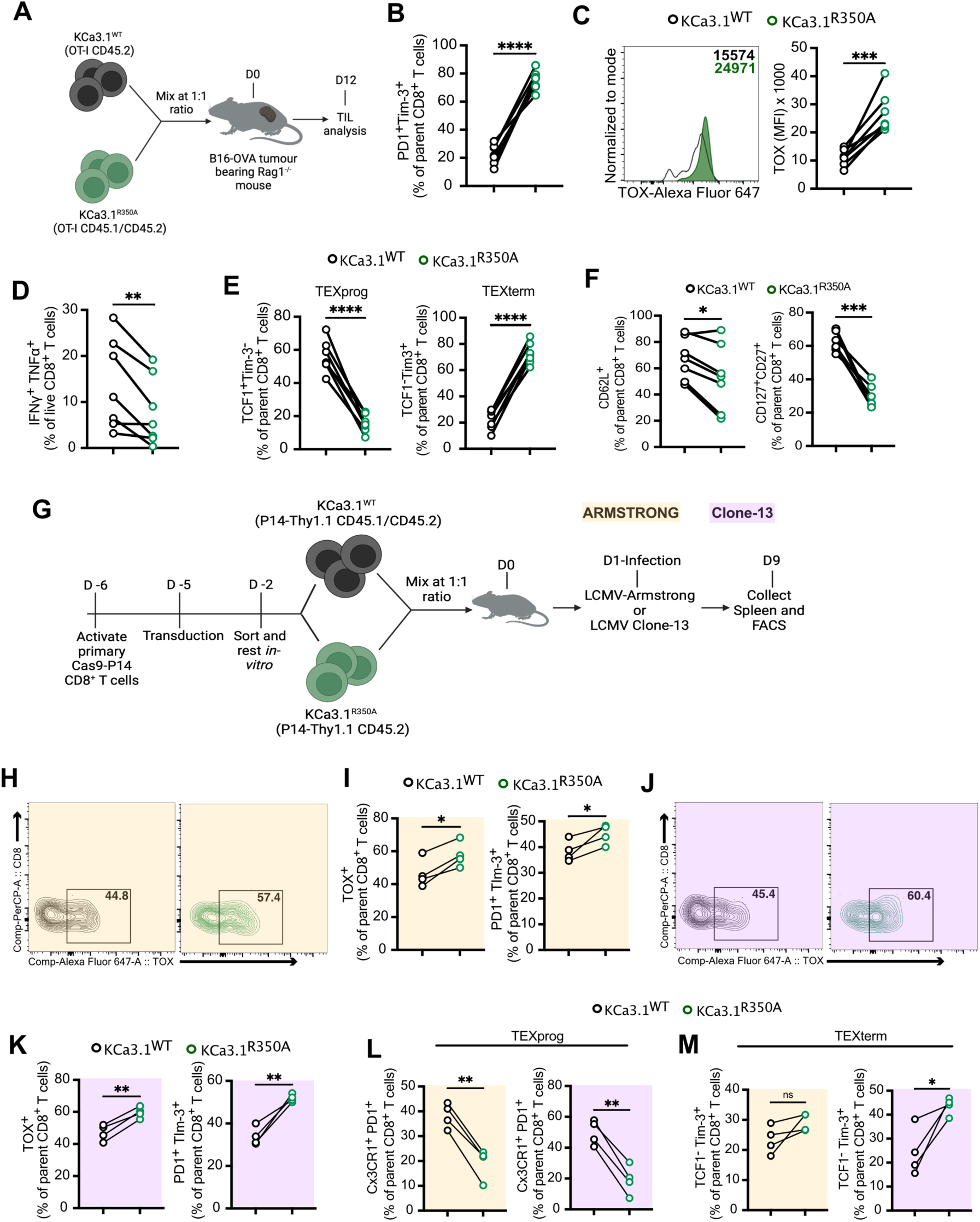
Ablation of KCa3.1 R350 methylation promotes T cell exhaustion in tumours and infection models. **a,** Schematic of experimental design to assess congenically distinct KCa3.1^WT^ and KCa3.1^R350A^ OT-I T cells that were mixed at a 1:1 ratio and transferred into B16-OVA tumour-bearing *Rag1*^-/-^ mice and analyzed day 12 post T cell transfer. **b-d** Frequency of Tim-3^+^PD1^+^ (b), histogram and quantification of TOX MFI (c), and frequency of IFNγ^+^TNFα^+^ (d) among KCa3.1^WT^ and KCa3.1^R350A^ OT-I TIL at D12 post T-cell transfer as in (a) (n=7). **e,** Frequency of TCF1^+^Tim-3^-^ TEXprog and TCF1^-^Tim-3^+^ TEXterm in KCa3.1^WT^ and KCa3.1^R350A^ OT-I TIL at D12 post T-cell transfer as in (a) (n=7). **f,** Expression of memory markers CD62L and CD127^+^CD27^+^ on OT-I T cells expressing KCa3.1^WT^ or KCa3.1^R350A^ isolated at D12 post transfer into B16-OVA tumour-bearing mice as in (a) (n=7). **g,** Schematic of experimental design to assess congenically distinct KCa3.1^WT^ and KCa3.1^R350A^ Thy1.1 P14 T cells that were mixed at a 1:1 ratio and transferred into WT mice infected with either LCMV-Armstrong or LCMV-Clone-13. Spleens were analyzed at D9 post infection. **h-i,** Expression of TOX (h) and frequency of TOX^+^ and PD1^+^Tim-3^+^ (i) in KCa3.1^WT^ and KCa3.1^R350A^ P14 CD8^+^ T cells from spleens at D9 post infection with LCMV-Armstrong (n=4). **j-k,** Expression of TOX (j) and frequency of TOX^+^ and PD1^+^Tim-3^+^ (k) in KCa3.1^WT^ and KCa3.1^R350A^ P14 CD8^+^ T cells from spleens at D9 post infection with LCMV-Clone-13 (p) (n=4). **l-m,** Frequency of CX3CR1^+^PD1^+^ TEXprog (l) and TCF1^-^Tim-3^+^ TEXterm (m) in KCa3.1^WT^ and KCa3.1^R350A^ P14 CD8^+^ T cells from spleens at D9 post infection as in (g) (n=4). Data are mean±s.e.m. Paired two-tailed Student’s t-test. *P < 0.05, **P < 0.01, ***P < 0.001, ****P < 0.0001.

T cell exhaustion is a prominent feature of chronic viral infection ^44^. We utilized the LCMV infection model in WT (CD45.2, Thy 1.2) mice into which CD45.1^+^CD45.2^+^ KCa.3.1^WT^ and CD45.2^+^ KCa.3.1^R350A^-expressing Thy1.1 (CD90.1^+^) P14^+^ T cells were injected at a 1:1 ratio into mice infected with either an acute Armstrong or a chronic Clone-13 strain. Spleens were analyzed on day 9 post-infection (Fig. 6G). The KCa.3.1^R350A^ population frequency was significantly reduced in Clone-13 infection, but no significant change was observed in acute infection (Extended Data Fig. 8J). KCa.3.1^R350A^-expressing T cells presented with increased TOX, PD1 and Tim-3 expression in both acute and chronic infections (Fig. 6H-K, Extended Data Fig. 8K). We also found that KCa.3.1^R350A^-expressing T cells harbored reduced frequencies of CX3CR1^+^PD1^+^ TEXprog cells (Fig. 6L) in both acute and chronic infection and increased Tim-3^+^Cx3CR1^+^PD1^+^ TEXeff cells (T exhausted effector-like cells) and TCF1^-^Tim-3^+^ TEXterm cells in chronic infection (Extended Data Fig. 8L and Fig. 6M). KCa.3.1^R350A^-expressing P14^+^ T cells displayed significantly increased KLRG1^+^CD127^-^ terminal effector-like cells and a concurrent decrease in KLRG1^-^ CD127^+^ memory precursor cells in chronic infection but not in acute infection (Extended Data Fig. 8M-O). Thus, ablation of R350 methylation drives T cells towards an exhaustion trajectory even at the early stages of infection. Together, these experiments support the idea that R350 methylation maintains optimal KCa3.1 function, ensuring optimal T cell activation and effector function in both tumours and viral infections.

## Acute Met supplementation promotes anti-tumour T cell function

CD8^+^ T cells are activated in both secondary lymphoid tissues and within tumours ^45,46^. While T cells are known to be primarily activated in draining lymph nodes (dLNs), a recent study showed that activation of tumour infiltrating T cells in tumour microenvironment is necessary to acquire an effector-like phenotype^47^. Our results in Fig. 1H and Extended Data Fig. 4G support the idea that Met is required for optimal TCR activation, and this requirement is independent of the prior activation status of the T cell. Thus, we hypothesized that upon tumour infiltration, CD8^+^ T cells experience antigen presentation in a Met-deficient microenvironment, leading to altered effector function. To evaluate Met levels in TILs, we performed mass spectrometric analysis of CD8^+^ TIL and dLN CD8^+^ T cells from B16 subcutaneous tumour-bearing mice at day 9. We observed a ∼50% reduction in intracellular Met in CD8^+^ TIL compared to that in dLN CD8^+^ T cells (Extended Data Fig. 9A, Supplementary table 2). We also observed a similar decrease in intracellular Met in CD8^+^ TIL isolated from primary human colorectal carcinomas compared with corresponding CD8^+^ T cells from PBMC (Extended Data Fig. 9B). To determine whether CD8^+^ TIL with reduced intracellular Met showed increased nuclear NFAT1, we sorted activated CD44^+^ CD8^+^ T cells from B16 tumours and respective dLN at day 9 post implantation and imaged for NFAT1. We observed increased levels of nuclear NFAT1 in CD44^+^CD8^+^ TIL compared to counterparts from dLN (Fig. 7A), suggesting that reduced Met availability in the TME may have led to increased NFAT1 activation in those T cells.

**Figure 7.**
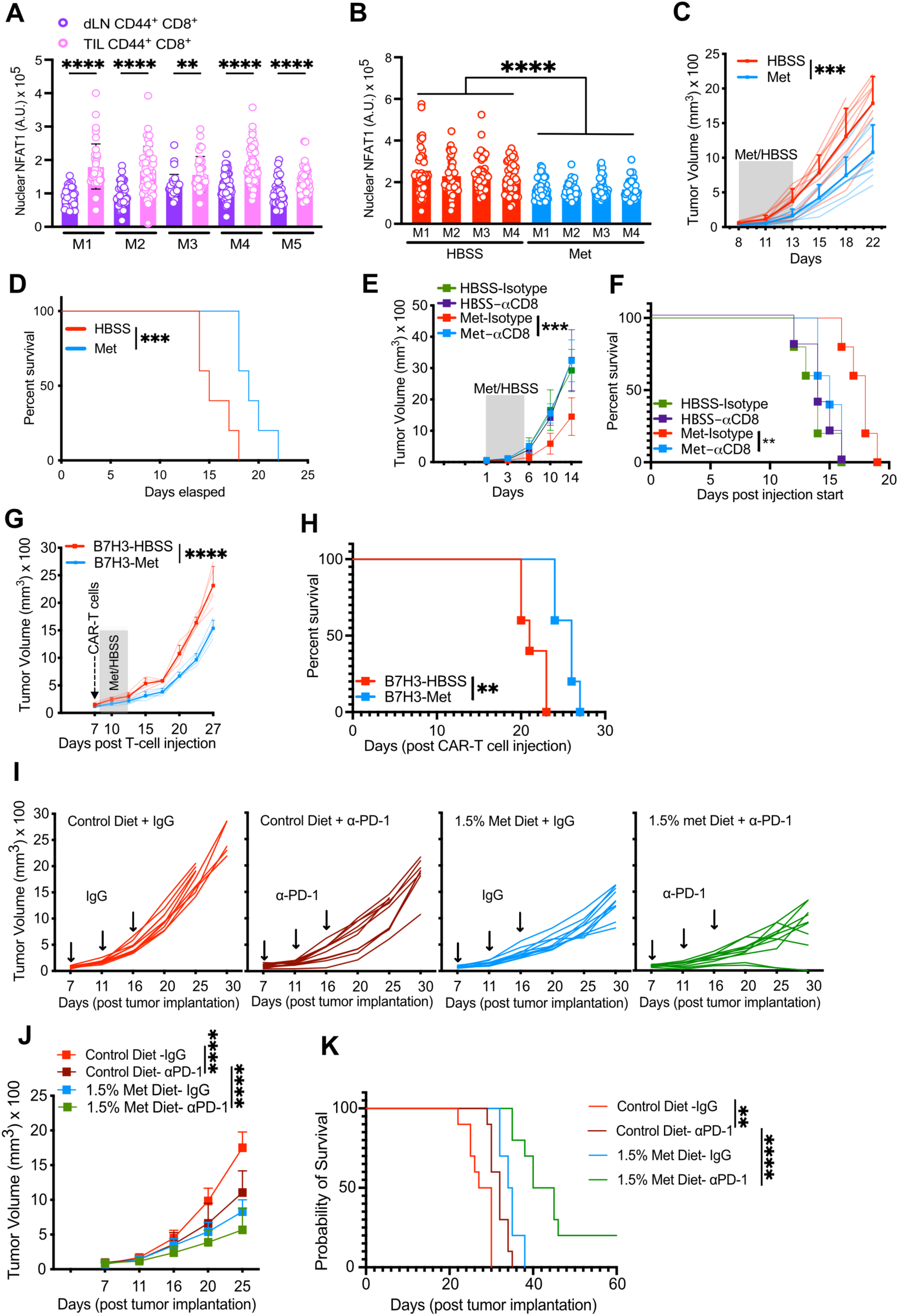
Acute methionine supplementation promotes CD8^+^ T cell-mediated tumour control and enhances ICB treatment. **a,** Nuclear NFAT1 quantification in CD44^+^CD8^+^ T cells isolated from B16 tumours and respective dLNs at D9 post implantation (each circle represents one cell, n=40 cells, n=5). **b,** Quantification of nuclear NFAT1 in CD44^+^ CD8^+^ T cells isolated from B16 tumours after 5 days of peri-tumoural supplementation of 50 μl HBSS or 61 μM Met per day (each circle represents one cell, n=40 cells, n=4). **c-d,** Tumour growth (c) and survival (d) of B16 tumour-bearing WT mice, supplemented either with HBSS or 61 μM Met peri-tumourally for five days as in (b) (n=10). **e-f,** B16 tumour growth (e) and survival (f) of WT mice, treated intraperitoneally with IgG isotype or anti-CD8 antibody at D1 and D3 and treated peri-tumourally with HBSS or Met from D7-D12 as in (b) (n=5, representative of two experiments). **g-h,** Tumour growth (g) and survival (h) of F420 tumour-bearing *Rag1*^-/-^ mice injected with B7-H3 CAR T-cells at D0 and peri-tumourally treated with HBSS or Met from D1-D6 as in (b) (n=5). **i-k,** Tumour growth (i, j) and survival (k) of MC38 tumour-bearing WT mice, treated either with anti-PD1 or IgG-isotype (shown by arrows in i) and fed with either control (1% Met) or Met-rich (1.5% Met) diet as shown in Extended Data Fig. 10H (n=10). Data are mean±s.e.m. Unpaired two-tailed Student’s t-test (a), 2-way ANOVA (c, e, g and j), and Mantel-Cox log rank test (d, f, h and k). Statistical analysis of (b) is described in methods. *P < 0.05, **P < 0.01, ***P < 0.001, ****P < 0.0001.

Continuous Met supplementation in tumours has been shown to promote anti-tumour T cell activity, associated with global epigenetic maintenance ^32^. Our findings, however, highlight an early role for Met in T cell fate, and we hypothesized that Met supplementation up to that of serum levels at the early stages of tumour development would increase T cell anti-tumour activity. We therefore injected 50 μl of 61μM Met peri-tumourally once per day for 5 days when tumour volumes were 70-100 mm^3^ (Extended Data Fig. 9C) and found that treatment decreased nuclear NFAT1 levels in CD44^+^ CD8^+^ TIL as compared to TIL from HBSS-injected tumours (Fig. 7B). Indeed, nuclear NFAT1 in TIL was similar to or lower than that in CD8^+^ T cells from dLN (Extended data Fig. 9D). Furthermore, acute Met supplementation delayed tumour growth in multiple murine tumour models, thereby improving overall survival (Fig. 7C, D and Extended Data Fig. 9E-G). Met supplementation resulted in decreased PD1 and Tim-3 surface expression in CD8^+^ TIL, along with increased CD62L^hi^CD44^hi^ Tcm and reduced CD62L^lo^CD44^hi^ Tem (Extended Data Fig. 9H and I) compared to TIL in HBSS-treated tumours. In contrast to immunocompetent WT mice, Met injected peri-tumourally in tumour-bearing NSG mice resulted in no difference in tumour growth (Extended Data Fig. 9J), implying a lymphocyte-mediated effect. Depletion with an anti-CD8 antibody (Extended Data Fig. 9K) in B16-OVA tumour-bearing mice abrogated the effect of Met supplementation on tumour control (Fig. 7E and F), supporting the conclusion that the observed Met supplementation-mediated tumour control is dependent on CD8^+^ T cells. We then employed a contralateral tumour model with B16-OVA implanted in one shoulder and in the opposite abdominal flank and injected Met peri-tumourally to the latter tumour. Met supplementation of the abdominal flank tumours impeded contralateral tumour growth (Extended Data Fig. 10A and B), further suggesting a T cell response. These results are consistent with the idea that acute Met supplementation promoted tumour control by CD8^+^ T cells. We also found that an 0.5% increase in dietary Met supplementation in B16 tumour-bearing mice showed better tumour growth control (Extended Data Fig. 10C), which is concordant with a recent study showing that Met supplementation in humans increased the effector function of CD8^+^ T cells ^32^. Met supplementation, therefore, enhances the anti-tumour function of CD8^+^ T cells.

## Met supplementation enhances CAR and ICB treatment

CAR-T cell treatment has been clinically successful in the treatment of hematological malignancies, but less so in solid tumours ^48,49^. In a solid tumour model in *Rag1*^-/-^ mice, we supplemented Met in the presence of transferred murine B7-H3 CAR-T cells into animals bearing a B7-H3^+^ F420 tumour^50^ (Extended Data Fig. 10D). Again, Met supplementation improved tumour control and animal survival in this model (Fig. 7G and H).

Immune checkpoint blockade (ICB) has been widely used either alone or in combination with cancer treatments^51^. To query whether Met supplementation might cooperate with ICB, we administered either IgG isotype or anti-PD1 antibody along with HBSS or Met supplementation in subcutaneous MC38 tumour-bearing mice (Extended Data Fig. 10E) and found that peritumoural Met supplementation with anti-PD1 treatment resulted in significantly reduced tumour growth and increased survival of these animals compared to controls (Extended Data Fig. 10F and G). As local Met supplementation may not be feasible in the clinical setting, we performed an ICB experiment with dietary supplementation of Met (Extended Data Fig. 10H). We observed that a diet containing 1.5% Met (0.5% increase in dietary Met) enhanced the efficacy of anti-PD-1, leading to significant control of tumour growth and survival of the mice, compared to that of anti-PD1 with control diet (Fig. 7I-K, Supplementary Table. 3). These results suggest the potential application of Met as a combinatorial treatment in the existing immunotherapies.

## Discussion

T cell exhaustion is generally understood to be a consequence of persistent antigen stimulation ^12,44^; however, situations exist in which antigen persistence does not result in exhaustion, such as in autoimmune T cells in Type I diabetes ^21^. It is therefore possible that in addition to chronic antigen exposure, nutrient limitation contributes to exhaustion ^52,53^. A recent study suggested that T cells can follow an exhaustion trajectory within hours of antigen recognition ^54^. Here, we propose that the state of T cell exhaustion is the result of a multifactorial pathway that involves the assimilation of different input signals, including nutrients, and is engaged as early as the initial recognition of cognate antigen.

Studies have shown that strong TCR signals result in hyperactivation and activation-induced cell death, whereas weak TCR signals result in dampened responses ^55^. NFAT family members are key regulators of T cell activation and interact with key partners to induce activation-associated genes including effector- and exhaustion-associated genes ^28^. Continuous or partner-less NFAT activation is associated with hyperactivation, anergy and exhaustion ^29,31,56^; therefore, NFAT activation must be tightly controlled for optimal T cell function, but little is known about how niche-specific nutrients regulate NFAT. We found that during rapid tumour proliferation in the early growth phase, CD8^+^ TIL display increased NFAT1 activation, which is regulated metabolically. Our data provide a novel mechanism of Met metabolism in the regulation of NFAT1 activation via post-translational modification of KCa3.1, without which CD8^+^ T cells progress towards exhaustion (Extended Data Fig. 11). We found that arginine methylation in KCa3.1 limits Ca^2+^-dependent NFAT1 signaling, ensuring optimal activation in tumours and infections. Met supplementation to the TME preserves anti-tumour immunity ^32^ and limits exhaustion (this study), a treatment that can be harnessed to improve patient prognosis in combination with ICB therapies. Our results emphasize the importance of non-histone protein methylation in the functional maintenance of T cells in diseases and describe a link between TCR signaling, metabolic regulation, and transcription factor activation. These results suggest a change in the current dogma of T cell exhaustion from a state driven only by persistent antigen exposure—acquired over a long period of time—to a dynamic state controlled by multiple pathways over the course of T cell activation and differentiation. Our study supports the idea that metabolism-dependent, rapid proteomic modifications that occur upon TCR engagement influence subsequent T cell fate and provides mechanistic insight into how niche-specific nutrient deprivation renders T cells dysfunctional

## Acknowledgements

We thank P. Fitzgerald, S. Sirasanagandla, M. Yang and L. Harris, for technical and administrative assistance; L. Long and H. Chi for providing retroviral vectors and Cas9-OTI mice; C. Guy for microscope imaging assistance; M. Morrison for organizing patient samples; the staff at the St. Jude Immunology Flow Cytometry Core Facility for cell sorting. Statistical calculations were performed using GraphPad Prism. We thank Dr. S. Pounds for the additional assistance with the statistical analysis of the data in Fig. 4A. Illustrations were created using BioRender software. This work was supported by National Institutes of Health grants AI123322 and CA231620 to D.R.G., American Lebanese Syrian Associated Charities (ALSAC) to G.K., and National Cancer Institute grant K99CA256262 to D.H.

## Author Contributions

P.S. conceived the project, designed, performed most experiments, interpreted results, and wrote the manuscript. A.G. contributed to CUT&RUN, ATAC-Seq, and RNA-Seq experiments. K.V. and E.B.R. helped in the development of the flow cytometry panel, experiments, and review of the manuscript. G.P. contributed to and processed the LC-MS/MS experiments. D.H. and G.K. assisted with CAR-T cell experiments. K.I. and M.B. performed MD simulations and analyses. S.P. and M.J.C. analyzed the CUT&RUN, ATAC-seq, and RNA-seq data. A. M. and J.P. performed and analyzed methyl-arginine immunoprecipitation and TMT-MS experiments. E.S.G provided patient tumour samples. D.R.G. supervised the experimental designs, interpreted the results, and wrote the manuscript.

---

**Extended Data Figure 1.**
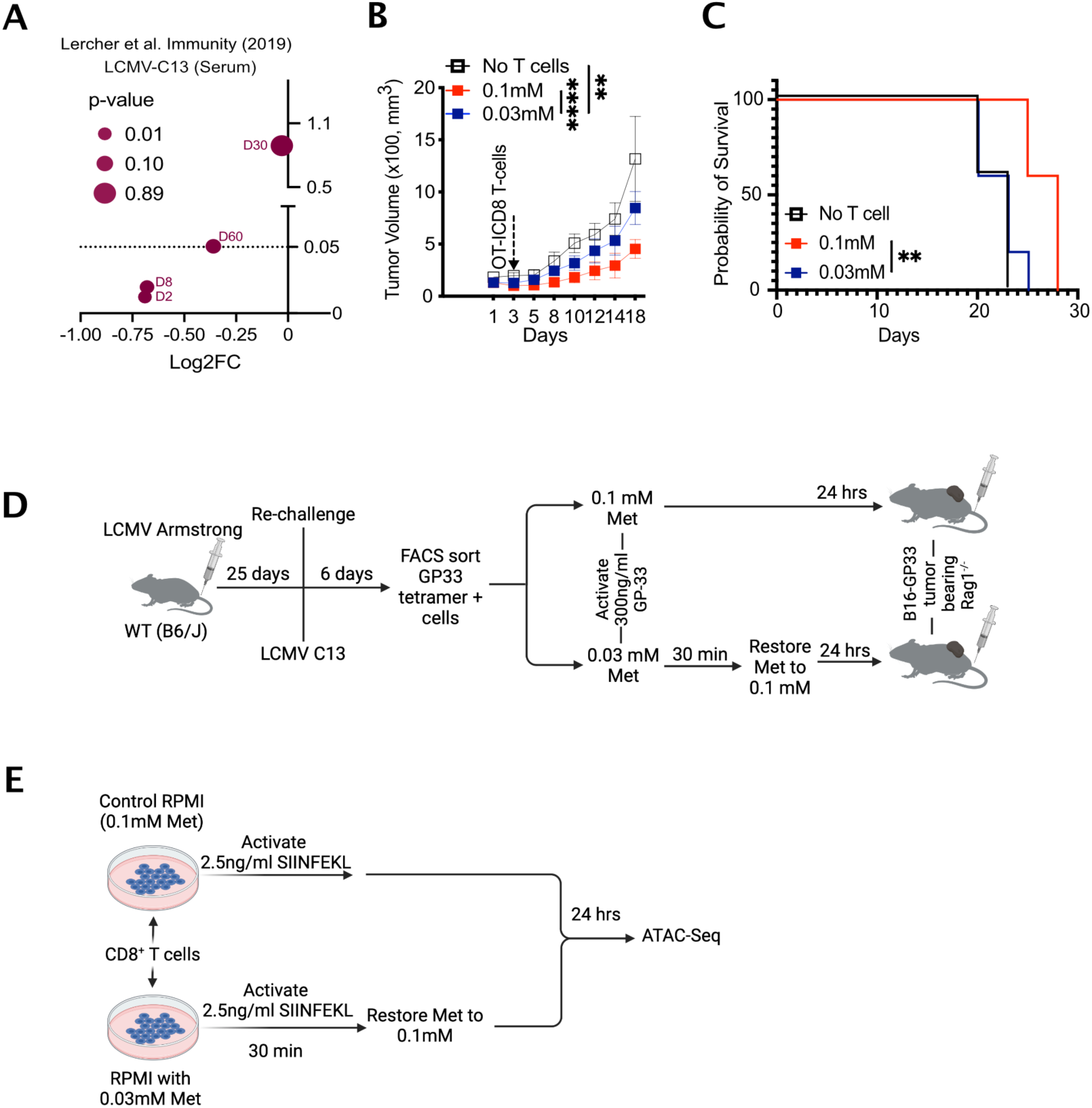
Methionine availability during TCR activation determines T cell fate. **a,** Log2FC in serum Met levels post LCMV-C13 chronic infection analyzed by Lercher et. al, *Immunity,* 2019 (ref. 7). **b-c,** Tumour growth (b) and survival (c) of B16-OVA tumour-bearing WT mice treated with or without OT-I CD8^+^ T cells activated in 0.1 or 0.03 mM Met for 30 min before restoring Met to 0.1 mM in 0.03 mM Met for 24 hrs, as described in Fig. 1d (n=5). **d,** Schematic of experimental design for generation of LCMV-GP33 specific memory T cells, which were then activated as described in (Fig. 1d) before injection into B16-OVA tumour-bearing *Rag1*^-/-^ mice. **e,** Schematic design for ATAC-Seq on OT-I T cell activation initially in 0.1 or 0.03 mM Met followed by restoration of Met to 0.1 mM for 24 hrs. Data are mean±s.e.m. 2-way ANOVA (b), and Mantel-Cox log rank test (c). **P < 0.01, ***P < 0.001.

**Extended Data Figure 2.**
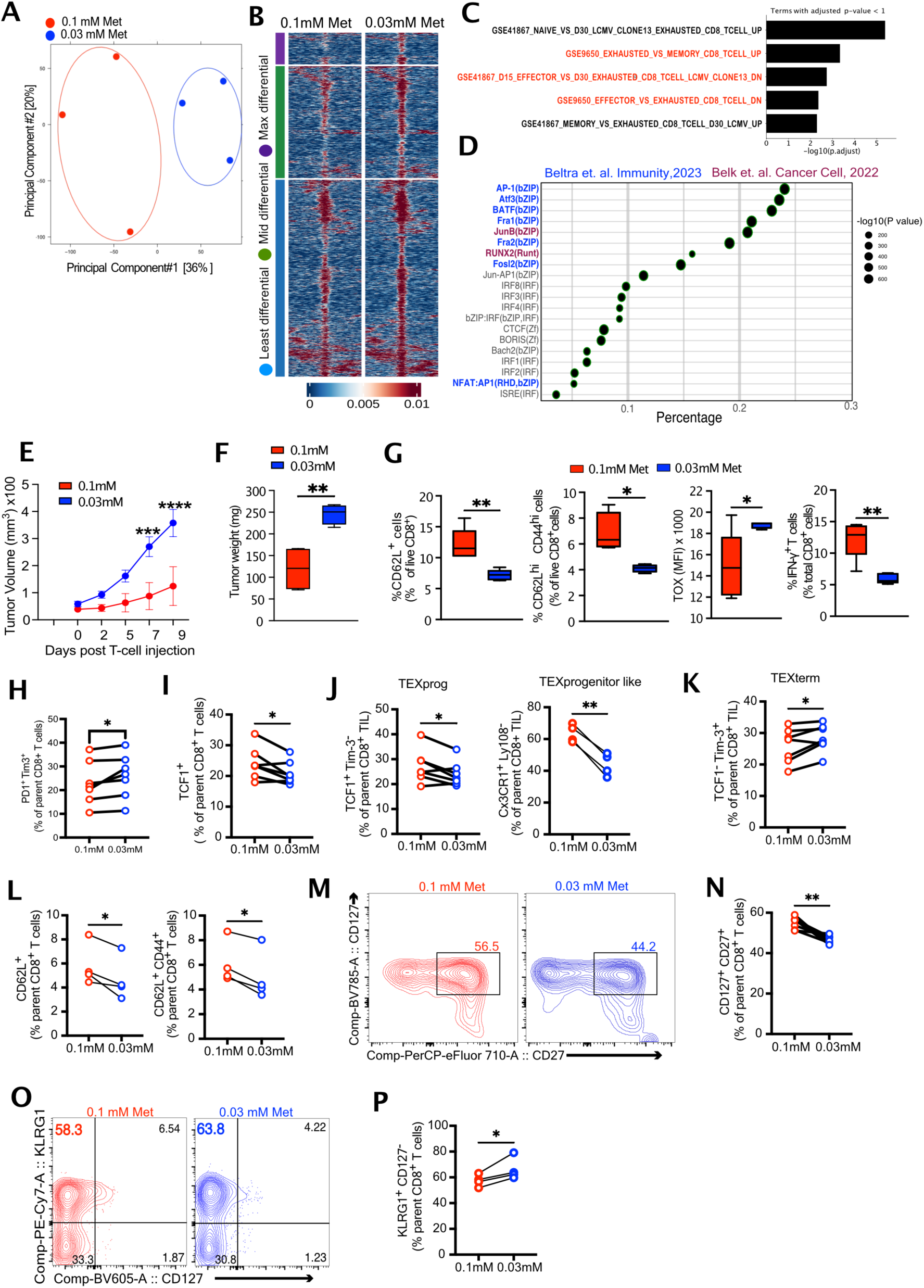
Reduced methionine during TCR activation promotes T cell exhaustion. **a,** PCA analysis of ATAC-Seq of OT-I T cells activated with 2.5ng/ml SIINFEKL in 0.1 mM and 0.03 mM Met and cultured in 0.1 mM Met for 24 hrs. n=3. **b,** Chromatin accessibility heatmap of T cells initially activated in 0.1 or 0.03 mM Met with each row representing peaks (p<0.05 and log2 fold-change >1.5) displayed over the span of 2kb window with peak as center (grouped from least to max differential region), analyzed from ATAC-Seq from (a). (n=3). **c,** Over-representation analysis of the DAR from (b) showing the top 5 genesets associated with exhaustion. **d,** Homer analysis of (a) showing top 20 motifs and their percent coverage. Colour corresponds to the common exhaustion-associated motifs also described in the indicated studies (blue and purple). **e-f,** Tumour growth (e) and weight (f) of B16-OVA tumour-bearing *Rag1*^-/-^ mice injected with OT-I T cells as in (Fig. 1d) (n=5). **g,** Frequency of CD62L^+^, Tcm (CD62L^hi^ CD44^hi^), TOX expression (MFI) and IFN-γ^+^ in OT-I CD8^+^ TIL isolated from B16-OVA tumour-bearing *Rag1*^-/-^ mice at D9 post-injection with OT-I T cells activated as in Fig. 1d (n=5). **h,** Frequency of PD1^+^ Tim-3^+^ on OT-I CD8^+^ T cells at D12 post transfer of OT-I T cells as in Fig. 2a (n=7). **i,** Frequency of TCF1^+^ on OT-I CD8^+^ T cells at D12 post transfer of OT-I T cells as in Fig. 2a (n=7). **j-l,** Frequency of TCF1^+^Tim-3^-^ TEXprog, CX3CR1^+^Ly108^+^ TEXprogenitor-like (j) and, TCF1^-^Tim-3^+^ TEXterm (k) CD62L^+^ T memory and CD62L^+^CD44^+^ T central memory cells (l) in OT-I CD8^+^ T cells as in Fig. 2a (n=7, n=4). **m-p,** Representative contour plot and quantification of CD127^+^CD27^+^ memory cells (m, n) and KLRG1^+^CD127^-^ terminal effector cells (o, p) in OT-I CD8^+^ TIL isolated at D12 post of cells activated as in Fig. 1d into B16-OVA tumour-bearing *Rag1*^-/-^ mice, as in Fig. 2a (n=4). Data are mean±s.e.m. Statistical analyses performed using 2-way ANOVA (e), unpaired two-tailed Student’s t-test (f, g) paired two-tailed Student’s t-test (h-p). *P < 0.05, **P < 0.01, ***P < 0.001.

**Extended Data Figure 3.**
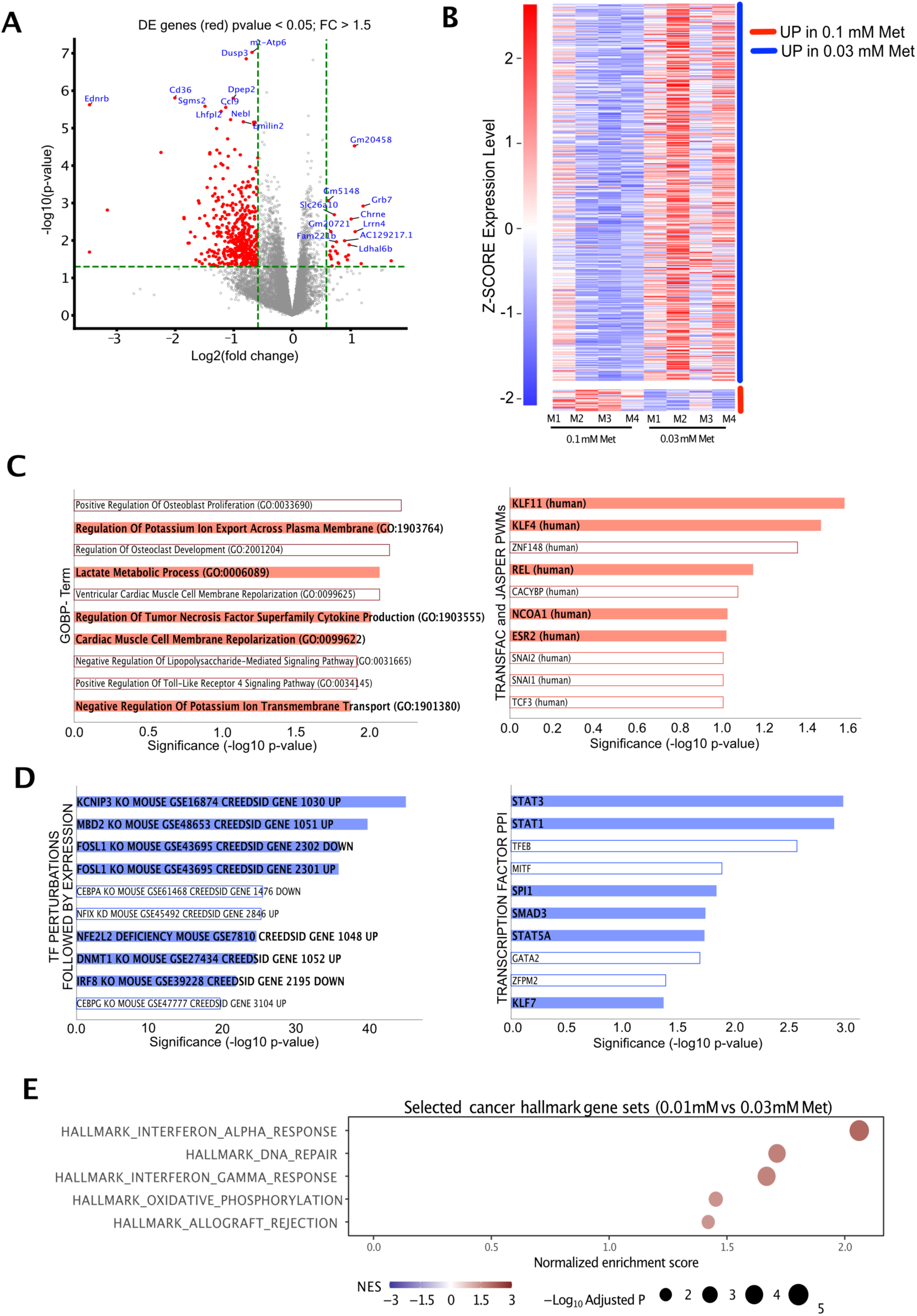
RNA-Seq analysis shows enrichment of exhaustion signature in low Met. **a,** Volcano plot and top ten differentially expressed genes (red) (0.1 mM vs 0.03 mM, p<0.05, FC>1.5) from OT-I CD8^+^ TIL isolated from mice injected with OT-I T cells, initially activated in 0.1 mM Met or 0.03 mM Met (as in Fig. 1d), at day 9 post injection into B16-OVA tumour-bearing mice (n=4). **b,** Heatmap of DE genes, which shows increased expression in cells initially activated in either 0.03 mM Met or 0.1 mM Met from the RNAseq in (a) (n=4). **c,** Over-representation analysis of DE genes with higher expression in TIL initially activated in 0.1 mM Met, showing enrichment of biological processes and transcription factors associated with T cell effector function and regulation. **d,** Over-representation analysis of DE genes with higher expression in TIL initially activated in 0.03 mM Met, showing enrichment of gene sets and transcription factors associated with promotion and maintenance of the exhausted T cell state. **e,** GSEA analysis of hallmark genesets as analyzed from RNA-Seq of OT-I TIL initially activated in 0.1 or 0.03 mM Met, isolated from B16-OVA tumours at D9 post transfer (n=4). RNA-Seq statistics were applied as described in methods.

**Extended Data Figure 4.**
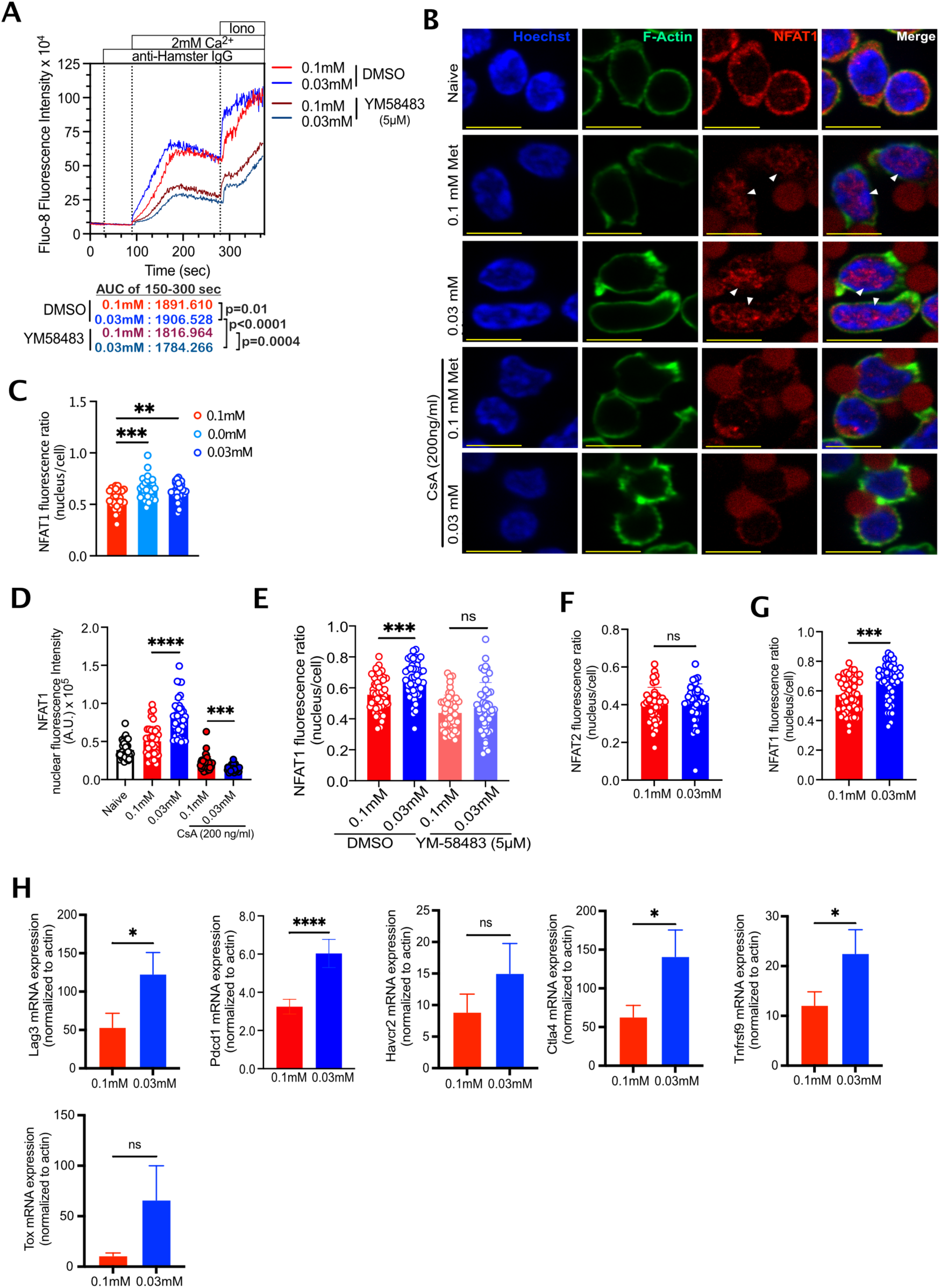
Met limitation results in increased TCR-mediated NFAT1 activation. **a,** Fluo-8 AM analysis of Ca^2+^ flux in CD8^+^ T cells activated with anti-CD3 and anti-CD28 and anti-hamster IgG crosslinking in Ca^2+^-free Ringer solution containing either 0.1 mM or 0.03 mM Met and treated for one hour with DMSO or 5μM YM58483. Area under the curve calculated from 150-300 sec (n=2). **b,** Un-edited image of Figure. 3b showing the unmasked, red anti-CD3/38 dynabeads which are masked grey in the main figure to highlight NFAT1 staining. **c,** NFAT1 intensity quantification (nuclear to total cell ratio) of CD8^+^ T cells activated in either 0.1 mM, 0.00 mM or 0.03 mM Met for 30 min with anti CD3/28 dynabeads and stained for NFAT1 (each circle represents one cell, n=40 cells). **d,** Quantification of nuclear NFAT1 from the experiment shown in Figure 3b. **e,** NFAT1 intensity quantification (nuclear to total cell ratio) of CD8^+^ T cells activated for 30 min by CD3/28 dynabeads in either 0.1 mM or 0.03 mM Met in the presence of DMSO or 5μM YM58483 (each circle represents one cell, n=40 cells). **f,** NFAT2 quantification (nuclear to total cell ratio) of CD8^+^ T cells, activated in 0.1 mM or 0.03 mM Met for 30 min with anti-CD3/28 dynabeads (each circle represents one cell, n=40 cells). **g,** NFAT1 quantification (nuclear to total cell ratio) of previously activated OT-I T cells, treated with anti-CD3/28 dynabeads for 30 min. T cells were initially activated with 2.5ng/ml SIINFEKL for 24 hrs in control medium and rested with 10 ng/ml mIL-7 for 48 hrs (each circle represents one cell, n=40). **h,** Normalized RNA expression quantified by quantitative-PCR performed 24 hrs post activation of OT-I CD8^+^ T cells activated with 2.5 ng/ml SIINFEKL in 0.1 mM or 0.03 mM Met for 30 min followed by 0.1 mM Met (n=6). Data are mean±s.e.m. Two-sample T-test (a) as described in the methods, unpaired two-tailed Student’s t-test (c-g) and one-tailed Student’s t-test (h). *P < 0.05, **P < 0.01, ***P < 0.001, ****P < 0.0001.

**Extended Data Figure 5.**
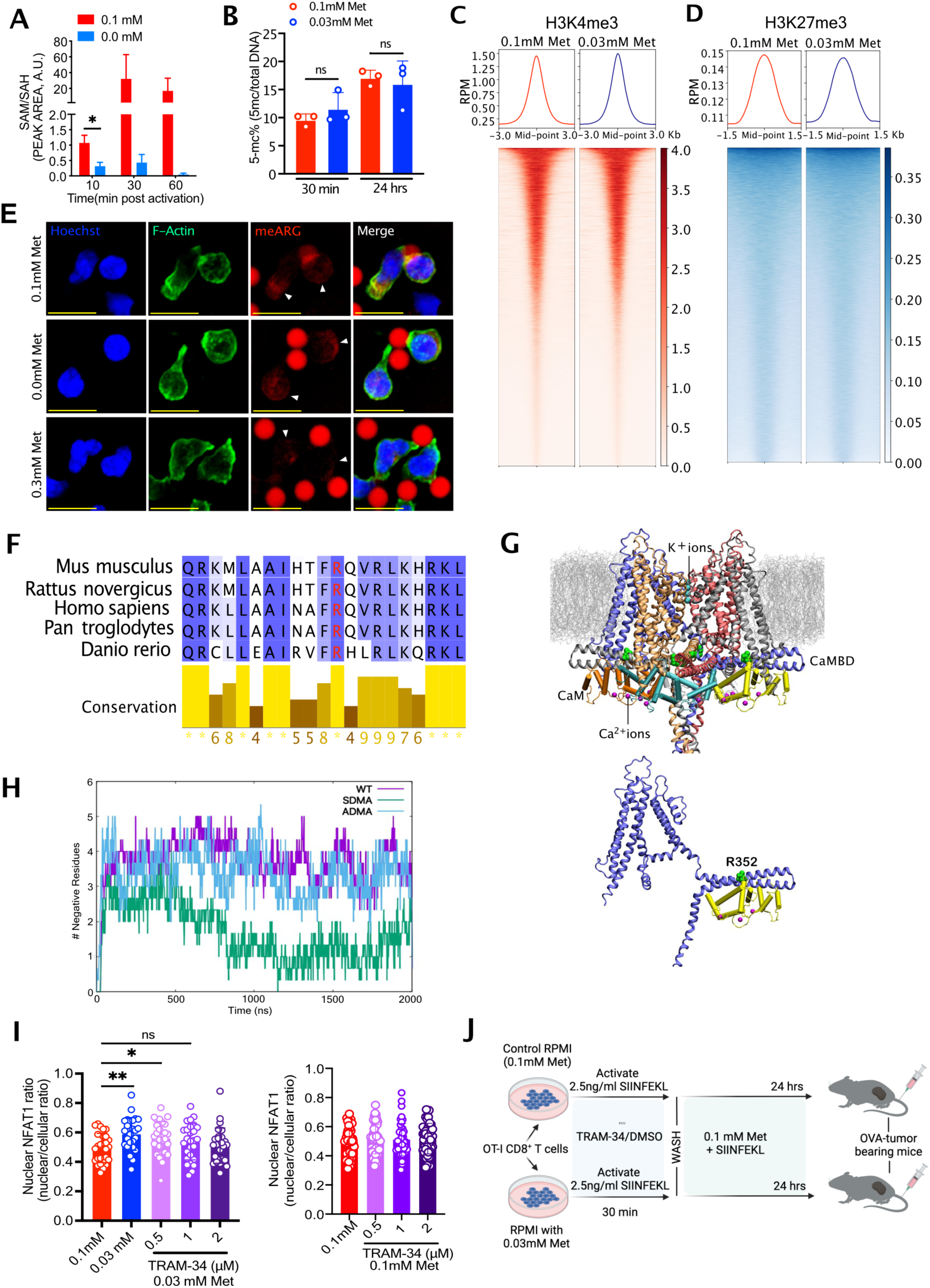
Initial Met limitation doesn’t affect DNA or histone methylation but alters KCa3.1 arginine methylation. **a,** Methylation potential (SAM/SAH) of 10 ng/ml SIINFEKL activated OT-I T cells in 0.1 mM or 0.0 mM Met at 10, 30 and 60 min (n=3). **b,** % 5-methylcytosine (%5-mc) of OT-I CD8^+^ T cells activated with 2.5 ng/ml SIINFEKL in 0.1 mM or 0.03 mM Met for 30 min, then 0.1 mM Met for 24 hr (n=3). **c-d,** Histograms (top) and heat-map (bottom) of H3K4me3 (c) and H3K27me3 (d) CUT&RUN reads (read count per million, normalized to background) in OT-I CD8^+^ T cells activated initially in 0.1 mM or 0.03 mM Met with restoration to 0.1 mM after 30 min and processed at 24 hrs post activation (n=3). **e,** Un-edited image of Figure. 4a showing the unmasked, red CD3/38 dynabeads which are masked grey in the main figure to highlight me-Arg staining. **f,** Conserved arginine (highlighted red) in KCa3.1 as analyzed by protein-BLAST in multiple species and visualized using Jalview. **g,** Tetrameric assembly of human KCa3.1-Calmodulin complex within the membrane (top). KCa3.1 is represented as ribbons and calmodulin quartet as cartoon with interacting Ca^2+^ ions as purple spheres. K^+^ ions are represented as cyan spheres. KCa3.1 monomer (blue) is shown in complex with calmodulin (yellow) along with demethylated arginine-352 (green) (bottom). **h,** MD simulations analysis: Illustration of total count of negatively charged residues within a 4Å radius of all four R352/SDMA/ADMA of huKCa3.1 the course of simulation time (averaged across all four monomers, three MD trials). **i,** NFAT1 quantification (nuclear to total cell ratio) in OT-I CD8^+^ T cells activated for 30 min with anti CD3/28 dynabeads in 0.03 mM Met (left) or 0.1 mM Met (right) in the presence of 0.5, 1 or 2 μM TRAM-34 (each circle represents one cell, n=45). **j,** Schematic design for OT-I T cell activation initially in 0.1 or 0.03 mM Met, treated with DMSO or TRAM-34 for 30 min, followed by washing and culturing in 0.1 mM Met+SIINFEKL (2.5ng/ml) for 24 hrs before injection into B16-OVA tumour-bearing *Rag1^-/-^* mice. Data are mean±s.e.m. Unpaired one-tailed Student’s t-test (a) or unpaired two-tailed Student’s t-test (b, i). *P < 0.05, **P < 0.01.

**Extended Data Figure 6.**
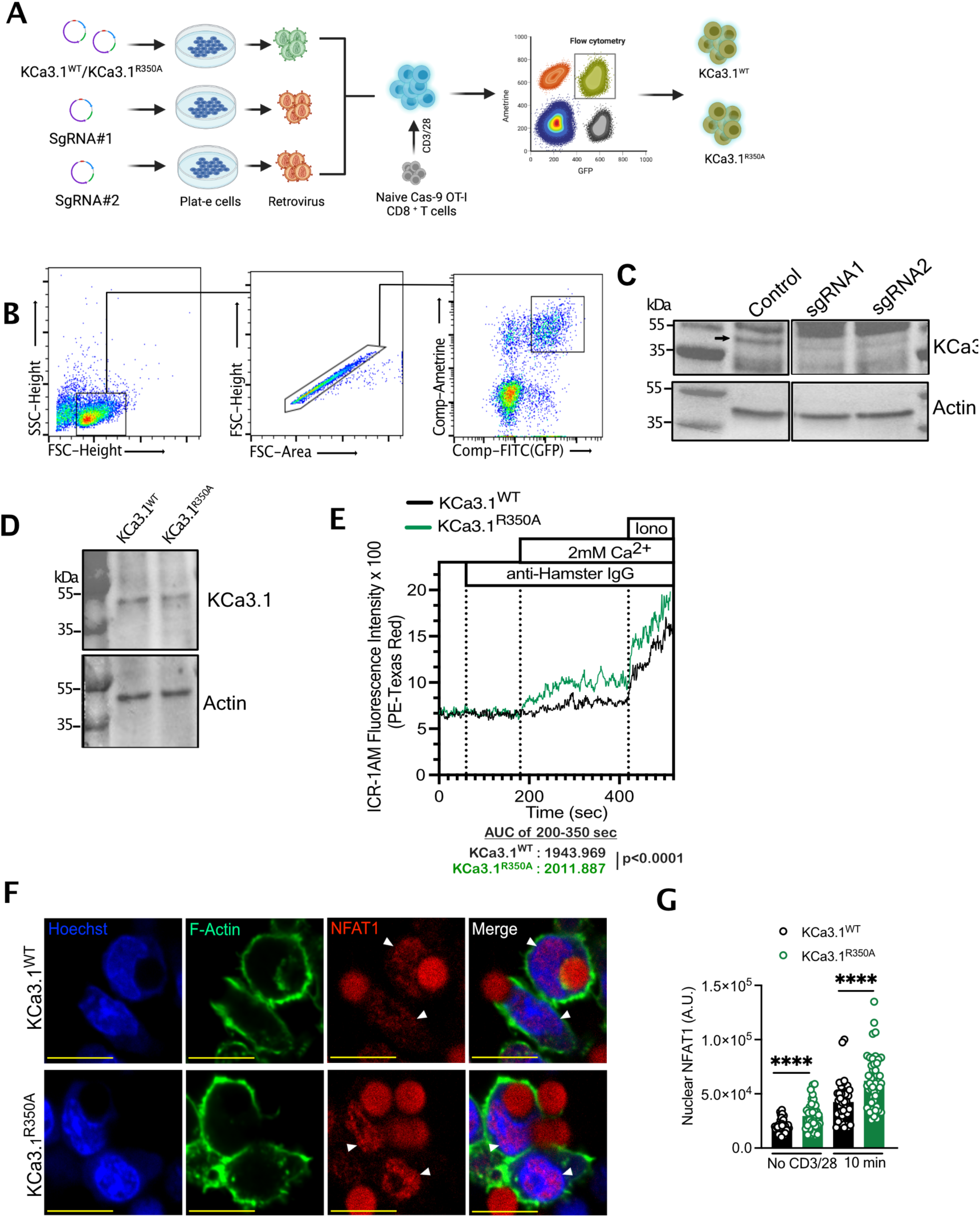
KCa3.1 R350 methylation regulates Ca^2+^-mediated NFAT1 activation. **a,** Schematic of experimental design of endogenous KCa3.1 knockdown with expression of KCa3.1^WT^ and KCa3.1^R350A^ in Cas9-OT-I or P14^+^ CD8^+^ T cells. **b,** Sorting strategy for *in vitro*-generated KCa3.1^WT^ and KCa3.1^R350A^ T cells; gate shows the cells that express WT or R350A KCa3.1 with endogenous KCa3.1 knockdown. **c,** Immunoblot showing knockdown of KCa3.1 with sgRNA1 and sgRNA2, compared to control. **d,** Immunoblot of KCa3.1 expression in KCa3.1^WT^- and KCa3.1^R350A^-expressing CD8^+^ T cells. **e,** ICR-1 AM analysis of Ca^2+^ flux in KCa3.1^WT^ and KCa3.1^R350A^ T cells activated with anti-CD3 and anti-CD28 by anti-hamster IgG crosslinking in Ca^2+^-free Ringer solution. Area under the curve is calculated from 200-350 sec. **f,** Un-edited image of Figure. 5b showing the unmasked CD3/38 dynabeads which are masked grey in the main figure to highlight NFAT1 staining. **g,** Nuclear NFAT1quantification of KCa3.1^WT^ and KCa3.1^R350A^, either control (no Dynabeads) or activated with anti-CD3/28 Dynabeads for 10 min (each circle represents one cell, n=45 cells). Data are mean±s.e.m. Two-sample T-test (a) as described in the methods and unpaired two-tailed Student’s t-test. ****P < 0.0001.

**Extended Data Figure 7.**
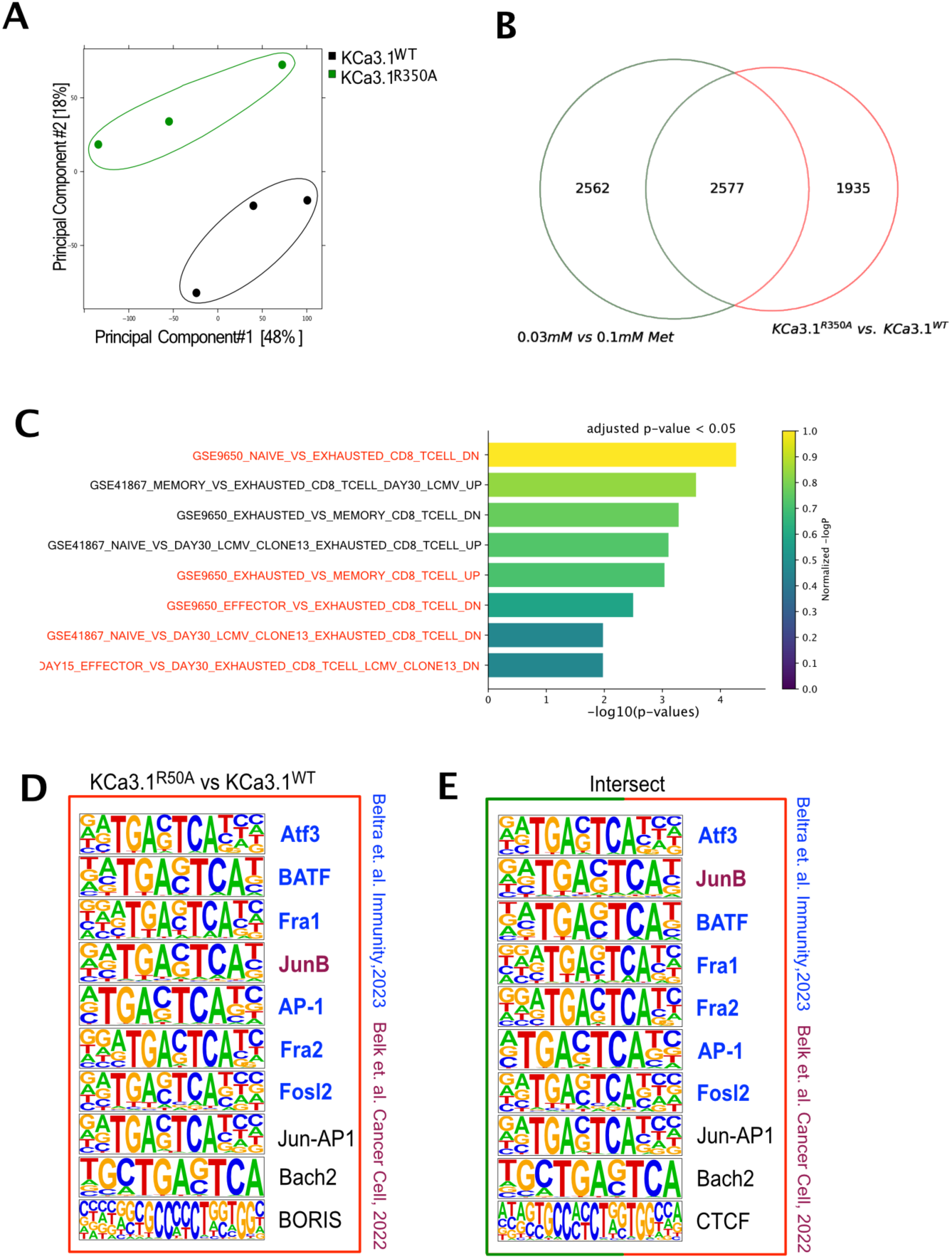
Ablation of KCa3.1 R350 methylation shows similar ATAC-Seq signature as of T cells activated in reduced Met. **a,** PCA analysis of ATAC-Seq of KCa3.1^WT^- and KCa3.1^R350A^-activated with 2.5 ng/ml SIINFEKL for 24 hrs. n=3. **b,** Common differentially accessible regions (DAR) (p<0.05, fold change>1.5) between ATAC-Seq of OT-I cells expressing KCa3.1^WT^ or KCa3.1^R350A^, or OT-I cells initially activated in 0.1 mM or 0.03 mM Met as in (a) and **Extended Data Fig. 2b. c,** Over-representation analysis of the common DAR from (b) showing the top 8 gene sets associated with exhaustion and activation. **d-e,** Top 10 motifs analyzed by HOMER in ATAC-Seq DAR’s from 24hrs activated OT-I CD8+ T cells expressing KCa3.1^R350A^ vs KCa3.1^WT^ (d), and from the common DAR’s from (b) (e). Colour corresponds to the common exhaustion-associated motifs also described in the indicated studies (blue and purple).

**Extended Data Figure 8.**
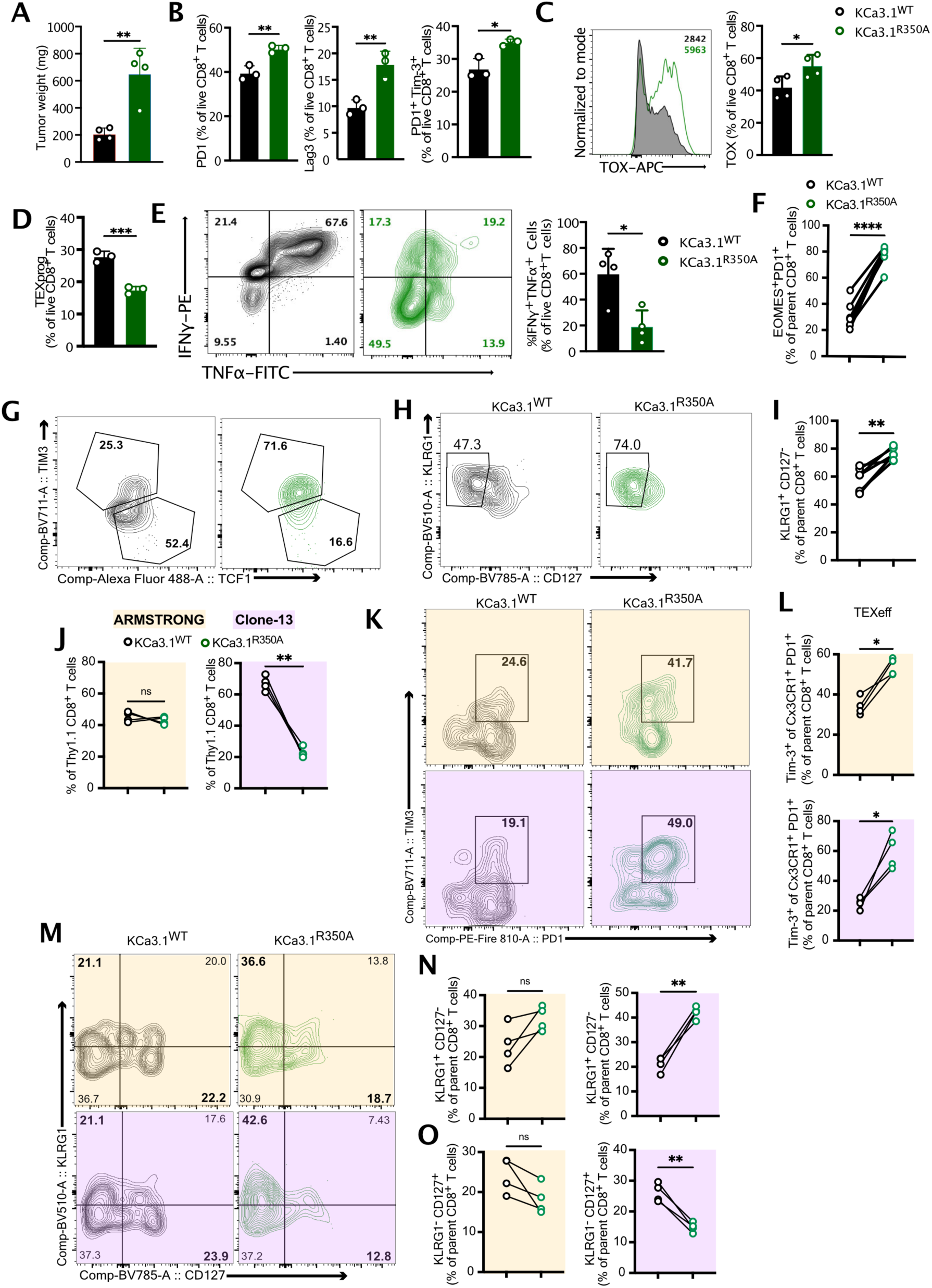
Ablation of KCa3.1 R350 methylation promotes T cell exhaustion *in-vivo*. **a,** Tumour weight of B16-OVA tumours isolated at D12 post transfer of OT-I T cells expressing KCa3.1^WT^ or KCa3.1^R350A^ (n=4). **b,** Surface expression of PD1^+^, Lag3^+^ and PD1^+^Tim-3^+^ in CD8^+^ TIL isolated at D12 post transfer into B16-OVA tumour-bearing mice of OT-I T cells expressing KCa3.1^WT^ or KCa3.1^R350A^ (n=4). **c,** Representative histogram of TOX expression and quantification in CD8^+^ TIL isolated at D12 post transfer of OT-I T cell expressing KCa3.1^WT^ or KCa3.1^R350A^ as (b) (n=4). **d,** Frequency of CD69^lo^Ly108^hi^ TEXprog in KCa3.1^WT^ and KCa3.1^R350A^ OT-I T cells at D12 post transfer (n=4). **e,** Representative contour plot (left) and quantification of IFNγ^+^TNFα^+^ (right) in OT-I T cells expressing KCa3.1^WT^ or KCa3.1^R350A^ at D12 post transfer as (b) (n=4). **f-g,** Expression of EOMES^+^PD1^+^ (f) and representative contour plot of expression of TCF1 and Tim-3 (g) on congenically marked OT-I T cells expressing KCa3.1^WT^ or KCa3.1^R350A^, mixed and transferred at a ratio of 1:1. TIL were isolated from B16-OVA tumours at D12 post T cell transfer (n=7). **h-i,** Representative contour plot (h) and quantification of KLRG1^+^CD127^-^ terminal effector cells (i) on congenically marked OT-I T cells expressing KCa3.1^WT^ or KCa3.1^R350A^ as in (f), isolated at D12 post transfer into B16-OVA tumour-bearing mice (n=7). **j,** Frequency of KCa3.1^WT^- and KCa3.1^R350A^-expressing P14 T cells in spleens, D9 post infection with either LCMV-Armstrong or LCMV-Clone-13 (n=4). **k,** Representative contour plot of PD1^+^Tim3^+^ KCa3.1^WT^- and KCa3.1^R350A^-expressing P14 T cells in spleens, D9 post infection with either LCMV-Armstrong or LCMV-Clone-13 (n=4). **j,** Frequency of Tim-3^+^Cx3CR1^+^PD1^+^ TEXeff KCa3.1^WT^- and KCa3.1^R350A^-expressing P14 T cells in spleens, D9 post infection with either LCMV-Armstrong or LCMV-Clone-13 (n=4). **m-o,** Representative contour plot (m) and quantification of KLRG1^+^CD127^-^ terminal effector cells (n) and KLRG1^-^CD127^+^ memory precursor cells (o) in spleens of animals injected with congenically marked P14 T cells expressing KCa3.1^WT^ or KCa3.1^R350A^, mixed 1:1, and infected with either LCMV-Armstrong or LCMV-Clone-13. Analysis on d9 post infection (n=4). Data are mean±s.e.m. Unpaired two-tailed Student’s t-test (a-e) and paired two-tailed Student’s t-test (f-n). *P < 0.05, **P < 0.01, ***P < 0.001, ****P < 0.0001.

**Extended Data Figure 9.**
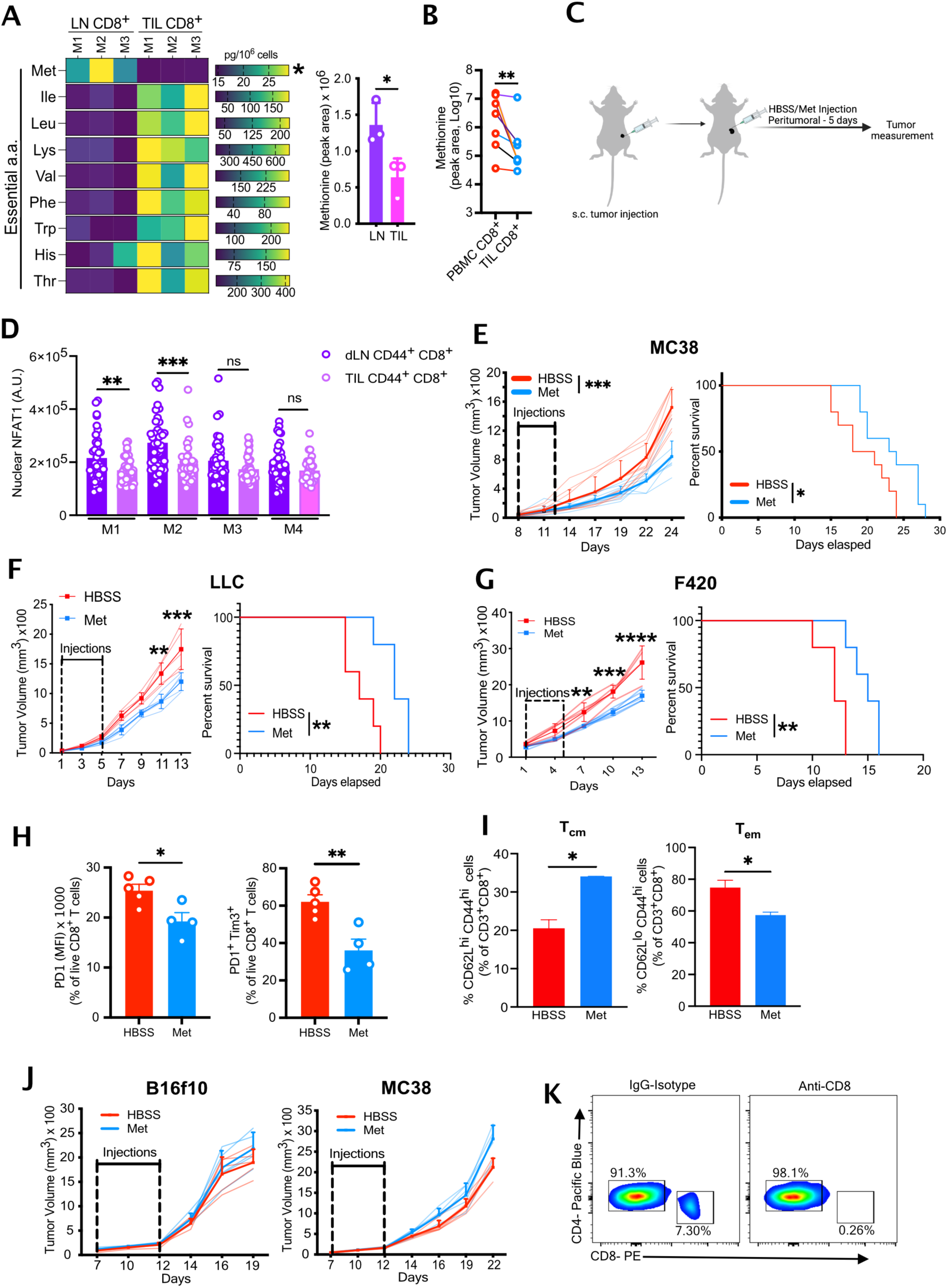
Acute Met supplementation promotes anti-tumour T cell differentiation and tumour control. **a,** Quantification of essential amino acids (left) and peak area of Met (right) in LN CD8^+^ T cells and CD8^+^ TIL, isolated at day 12 post tumour implantation (n=3). **b,** Quantification of intracellular Met in CD8^+^ PBMC and CD8^+^ TIL isolated from patient primary colorectal tumours (n= 8). **c,** Schematic of experimental design of acute HBSS/Met peritumoural treatment in sub-cutaneous tumours. **d,** Nuclear NFAT1 quantification of CD44^+^ CD8^+^ T cells isolated from B16 subcutaneous tumours and dLN, 5 days post daily peri-tumoural supplementation of 50 ml 61μM Met (M=1 mouse, each circle represents one cell, n=40 cells). **e-g,** Tumour growth (left) and survival (right) of WT mice with sub-cutaneous MC38 colorectal tumours (e) (n=10), Lewis-lung carcinoma (LLC) tumours (f) (representative of two experiments of n=5), and F420 osteosarcoma tumours (g) (n=5), treated with either HBSS or 50 ml 61μM Met peri-tumourally daily for 5 days. **h-i,** Expression of PD1, PD1^+^Tim-3^+^ (h), Tcm (CD62L^hi^Cd44^hi^) and Tem (CD62L^lo^CD44^hi^) (i) on CD8^+^ TIL isolated from B16 tumour-bearing mice 2 days after 5 daily peri-tumoural injection with HBSS or Met (n=5-4). **j,** Tumour growth of B16f10 and MC38 tumours in NSG mice treated peri-tumourally with 50 μl HBSS or 50 μl 61μM Met daily for 5 days (n=5). **k,** Representative flow cytometry plot of blood showing CD8^+^ T cell depletion in anti-CD8 antibody treated mice compared to IgG-Isotype-treated mice (n=4). Data are mean±s.e.m. Unpaired (a, d, h and i) and paired (b) two-tailed Student’s t-test, two-way ANOVA (e-g (left) and j), Mantel-Cox log rank test (e-f (right)). *P < 0.05, **P < 0.01, ***P < 0.001, ****P < 0.0001.

**Extended Data Figure 10.**
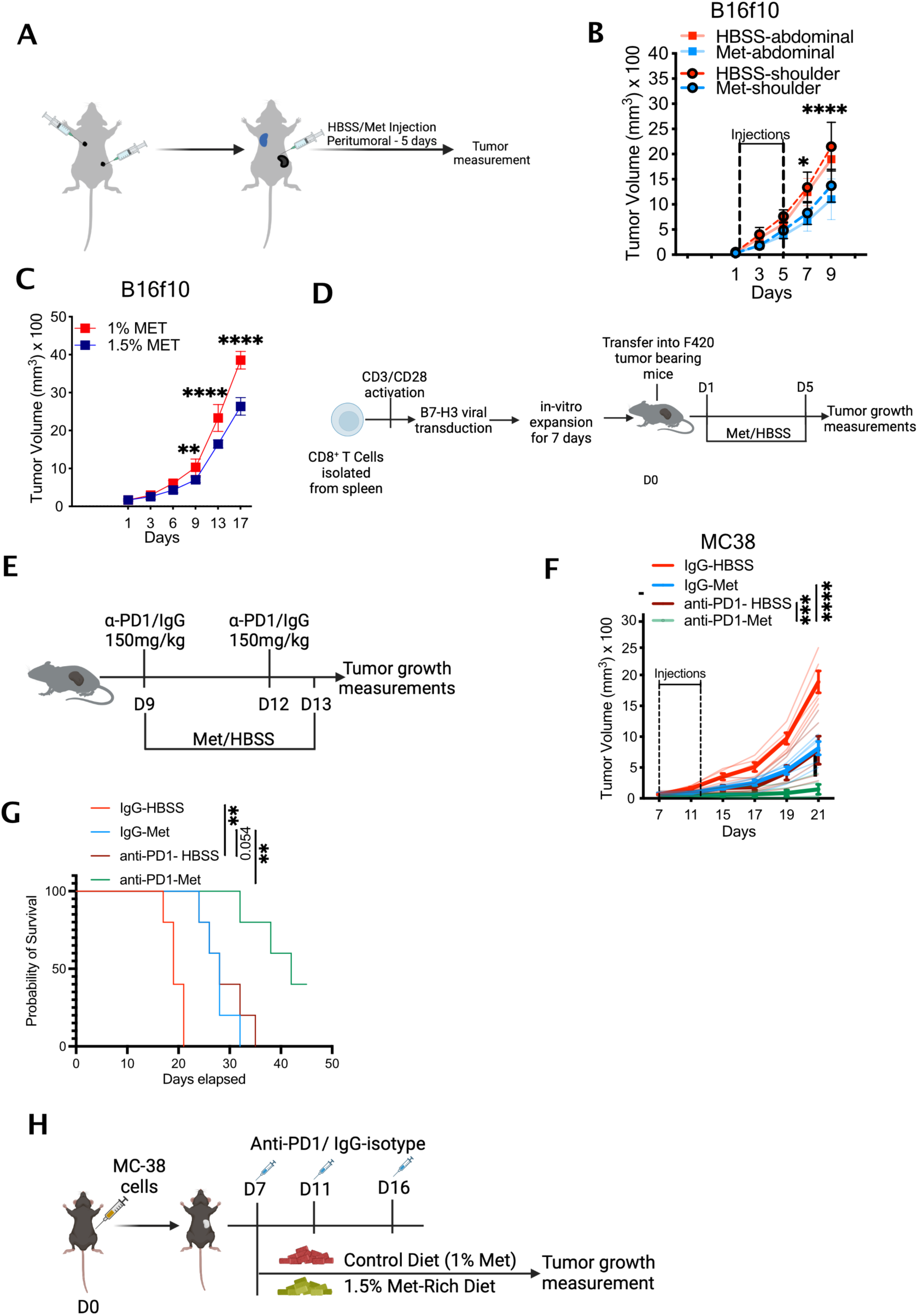
Acute Met supplementation promote T cell mediated tumour control and enhances tumour immunotherapies. **a,** Schematic of experimental design of contralateral tumour experiment. **b,** Growth of B16f10 tumours implanted on left shoulder (dashed lines) and right flank (solid lines) following peri-tumoural injection of flank tumour with 50 μl HBSS or 61μM Met daily for 5 days (n=5, representative of two experiments). **c,** B16 tumour growth in WT mice fed with either 1 % Met chow or 1.5 % Met chow for 7 days post tumour implantation (n=5). **d,** Schematic of experimental design to assess the effect of HBSS or Met supplementation on CAR-T cell therapy in murine solid tumour model. **e,** Schematic of design to assess tumour growth with anti-PD1 treatment and with either peri-tumoural HBSS or Met supplementation. **f-g,** Tumour growth (f) and survival (g) of MC38 tumour-bearing WT mice, treated either with anti-PD1 or IgG-isotype and supplemented with HBSS or Met peri-tumourally for five days as in (e) (n=5, representative of two experiments). **h,** Schematic of experimental design to assess tumour growth with anti-PD1 treatment on sub-cutaneous tumour-bearing mouse, supplemented with either control diet or Met-rich diet. Data are mean±s.e.m. 2-way ANOVA (b, c, and f), and Mantel-Cox log rank test (g). *P < 0.05, **P < 0.01, ***P < 0.001, ****P < 0.0001.

**Extended Data Fig. 11:**
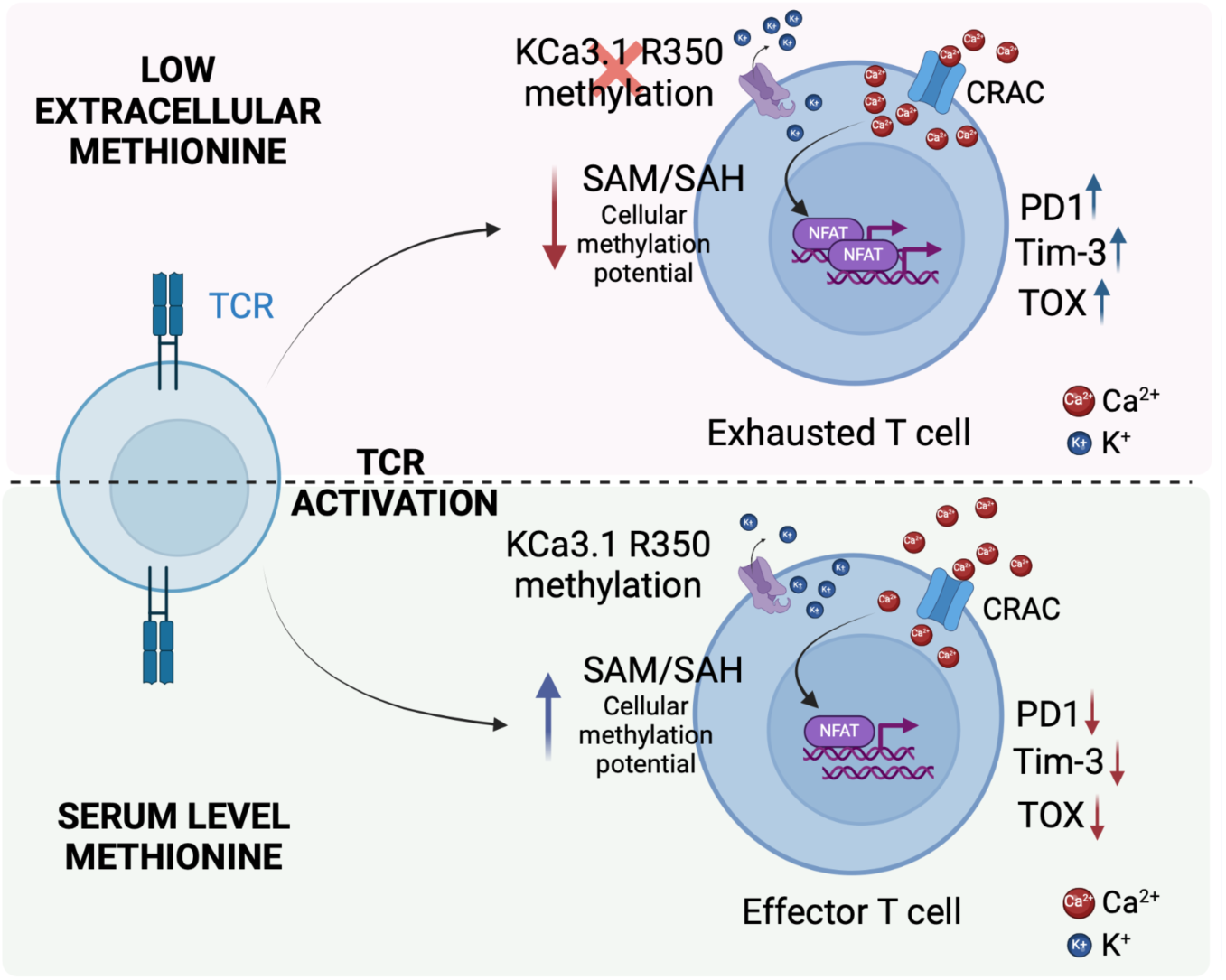
Methionine metabolism regulates TCR-mediated Ca^2+^ flux and T cell fate. Diagram illustrating the impact of methionine metabolism on TCR-dependent methylation of KCa3.1, leading to regulation of Ca^2+^ flux and subsequent activation of NFAT1. Low extracellular Met levels lead to decreased methylation potential, reducing KCa3.1 R350 methylation. This results in increased Ca^2+^ flux and downstream NFAT1 activation and consequent T cell hyperactivation and exhaustion.

## Materials and Methods

### Reagents

RPMI 1640 (cat. 11875093, Gibco) and methionine-free RPMI 1640 (cat. A1451701; Gibco) supplemented with 10% dialyzed FCS (Atlanta Biologicals), sodium pyruvate (cat. 11360070, Gibco), NEAA (11140050, Gibco), glutamine (cat. 25030081, Thermo-Fisher), and penicillin-streptomycin cocktail (cat. 15070063, Thermo-Fisher) and stored at 4°C. Methionine (cat. M5308, Sigma-Aldrich) was reconstituted in PBS (0.1 M) and stored at 4°C. SIINFEKL peptide (cat. 60193, AnaSpec) peptide was reconstituted in PBS at 0.5 mg/ml and stored at −20°C. TRAM-34 (cat. HY-13519, MedChem Express) was reconstituted at 1 M with DMSO. Cyclosporin A (CsA) (cat. 30024, Sigma-Aldrich) was reconstituted to a concentration of 50 mg/ml in DMSO. YM-58483 (cat. 3939, Tocris) was reconstituted at 20 mM stock in DMSO. All reconstituted reagents were stored at −20°C and aliquoted to avoid repeated thawing.

### Mice

All mouse studies were conducted in accordance with protocols approved by the St. Jude Children’s Research Hospital Committee on Care and Use of Animals, and in compliance with all relevant ethical guidelines. All mice were kept under a 12h-12h light-dark cycle in a specific-pathogen-free (SPF) facility at the institute’s Animal Resource Center. Age-and sex-matched mice were randomly assigned to the experimental and control groups. Cas9-OT/P14^+^-I (Cas9 Knock-in with OVA-specific transgenic TCR (OT-I) or LCMV-GP33 specific TCR(p14)), C57BL/6J (WT), OT-I, and *Rag1*^−/−^ mice were bred in-house. The investigators were not blinded to the experiments and outcome assessments.

### Cell lines

B16F10, B16F10-Ova (B16-Ova) melanoma cells were provided by Dr. Hongbo Chi. MC38 and MC38-Ova colon carcinoma cell lines were provided by Dr. Dario Vignali. The F420 sarcoma cell line was provided by Dr. Jason T Yustein. HEK293T and Lewis lung carcinoma (LLC) cells were purchased from the ATCC. Plat-e cells used for retroviral packaging were purchased from Cell Biolabs (cat. RV-101). All the cell lines used were tested and confirmed to be mycoplasma-negative but not independently authenticated.

### Murine and Human CD8^+^ T cell isolation and stimulation

Murine CD8^+^ T cells were isolated from the spleen and lymph nodes (LN) by physical disruption through a 70μ strainer before treatment with erythrocyte-lysis buffer. Cells were subjected to CD8^+^ T cell enrichment using the EasySep mouse CD8^+^ T cell isolation kit (cat. 19853, StemCell), according to the manufacturer’s protocol. Isolated T cells were then either activated with SIINFEKL peptide or mouse CD3/28 Dynabeads (cat. 11456D, Thermo-Fisher) in RPMI medium, as described above. Human CD8^+^ T cells were isolated from apheresis rings from the St Jude Blood Donor Center. Blood obtained from an apheresis ring was mixed at a 1:1 ratio with PBS containing 2% FBS and carefully layered onto Lymphoprep (cat. 07801, StemCell) solution. The samples were then centrifuged at 800g for 30 min at room temperature without braking. The peripheral blood mononuclear cell (PBMC) layer was aspirated and washed twice in PBS with 2% FBS before enrichment of CD8^+^ T cells using the Human CD8^+^ T Cell Isolation Kit (cat. 17953, StemCell), according to the manufacturer’s protocol. CD8^+^ T cells were activated using human CD3/28 Dynabeads (cat. 11131D, Thermo Fisher Scientific) in RPMI medium. Apheresis rings were collected with written consent from the donors for research upon review and approval by the Institutional Review Board at St Jude Children’s Research Hospital.

### Ca^2+^ Flux measurement and analysis

Isolated T cells were labelled with 3 μM Fluo-8 AM (AAT Bioquest, cat. 21080) or 5 μM ICR-1AM (Ion Biosciences) with 0.04% pluronic F-127 (Thermo-Fisher, cat. P6867) for 2 hrs in HHBS at 37 °C and washed twice with Ca^2+^-free Krebs-Ringer’s solution (Thermo Fisher Scientific, cat. no. J67839-AP). The cells were then resuspended in Ca^2+^-free Krebs-Ringer’s solution and incubated with anti-CD3ε (4 μg/ml; 2C11, cat. BE0001, Bio-X-Cell) and anti-CD28 (4 μg/ml; 37.51, cat. BE0015, Bio-X-Cell) on ice for 10 min. 0.1 mM Met or 0.03 mM Met was added to the samples and the baseline was recorded before the addition of hamster-IgG for crosslinking. Ca^2+^ influx was measured after the addition of 2mM Ca^2+^, 1uM ionomycin was added as a positive control for Ca^2+^ flux. All measurements were recorded on either Cytek Aurora (SpectroFlo) or BD-Fortessa flow cytometer.

To analyze temporal changes in fluorescence intensity, we binned the time variable into 10-second intervals to calculate the mean log2-transformed fluorescence intensity within each 10-second bin for each sample. We calculated the AUC across the entire time series, as well as within a focused time window (x–y seconds) using the trapezoidal rule, which sums the area under each segment formed by the binned time points. This approach captures cumulative fluorescence intensity over time, providing a metric for comparing samples. To evaluate statistical significance between groups, we performed two-sample t-tests on the log2-transformed fluorescence values within the specified time window (x–y seconds).

### Quantitative PCR

Isolated T cells were activated with 2.5ng/ml SIINFEKL in 0.1 mM Met or 0.03 mM Met for 30 min after which Met in 0.03 mM was restored to 0.1 mM. T cells were then further cultured for 24 hrs. Cells were washed twice with PBS and RNA was isolated using Direct-zol RNA miniprep Kit (Zymo research, cat. no. R2050). RNA was quantified and processed for cDNA synthesis using Reverse Transcriptase (Thermo Fisher Scientific, cat. no. 28025-013). cDNA was then used to perform quantitative PCR with the specific primers (Table S2) using SYBR green PCR master mix (Thermo Fisher Scientific, cat. no. 4309155). PCR reaction was run and quantified by QuantStudio 7 flex.

### LCMV infection

LCMV Armstrong and Clone-13 was obtained from Dr. Shaabani, Scripps Research Institute. The virus was propagated in BHK21(baby hamster kidney) cells as described^1^ and stored at −80 degree Celsius. For experiments, 2×10^6^ pfu of both Armstrong and Clone-13 was used at 2×10^6^ pfu/animal. The virus was injected intravenously, and the infected animals were maintained under a 12h-12h light-dark cycle in a SPF, Biosafety Level-2 facility in the institute’s Animal Resource Center.

### Cloning and virus production

*Kcnn4* gRNA’s were designed using the Broad Institute platform (https://portals.broadinstitute.org/gpp/public/analysis-tools/sgrna-design) and cloned into the ametrine-expressing retroviral vector LMPd-gRNA-mPGK-Ametrine (sgRNA1:CACCGGGCAGGCTGTCAATGCCACG, AAACCGTGGCATTGACAGCCTGCCC; sgRNA2:CACCGTGTGGGGCAAGATTGTCTGC, AAACGCAGACAATCTTGCCCCACAC). KCa3.1^WT^ and KCa3.1^R350A^ cDNA were synthesized by Genscript; the latter contained an alanine substitution at R350. Both KCa3.1^WT^ and KCa3.1^R350A^ had substitutions at the PAM recognition sites of the gRNAs (Table S3). Each cDNA was subcloned into the pMIGII. Plat-e cells were co-transfected with the cloned vectors and pCL-eco (cat. 12371, Addgene) in Opti-MEM (cat. 31985070, Thermo-Fisher) with Trans-IT-293 transfection reagent (cat. MIR 2705, Mirus Bio) and cultured at 37°C for 12-15 hrs prior to changing the medium and culturing for an additional 24 hrs at 37°C, followed by 24 hrs at 32°C, 5% CO_2_. The viral supernatant was harvested and spun at 1800rpm at RT to remove the cell debris. Viral supernatant was either used immediately or stored at −80°C till further use.

### Viral transduction

Naive Cas9-OT-I CD8^+^ cells were isolated from the spleen and peripheral LNs of Cas9-OT-I mice using a Magnisort naïve CD8^+^ T cell isolation kit according to the manufacturer’s instructions (cat. 8804-6825-74, Thermo-Fisher). Purified CD8^+^ T cells were activated *in vitro* for 20 h with plate-bound anti-CD3ε (5 μg/ml; 2C11, cat. BE0001, Bio-X-Cell) and anti-CD28 (5 μg/ml; 37.51, cat. BE0015, Bio-X-Cell) antibodies. Viral transduction was performed by spinfection at 900 *g* at 25 °C for 3 h with 10 μg/ml polybrene (Sigma-Aldrich), followed by resting for 3 h at 37 °C in 5% CO_2_. Cells were washed and cultured in medium supplemented with mouse rIL-2 (20 U/ml; cat. 212-12, Peprotech) for 4 days. GFP-Ametrine double-positive cells were sorted using the Reflection cell sorter (iCyt) or the MoFlo XDP cell sorter and rested for 24 hrs in medium containing rIL-2 (10U/ml, cat. 212-12 Peprotech) before activation or adoptive transfer into the animals.

### Flow cytometry

Antibodies from BioLegend included Pacific Blue anti-mouse Ly108 (330-AJ, 134608), BV510 anti-mouse KLRG1 (2F1, 138421), BV570 anti-mouse CD62L (MEL-14, 104433), BV711 anti-mouse Tim-3 (RMT3-23, 119727), BV785 anti-mouse CD127 (A7R34, 135037), PE-Cy5 anti-mouse Granzyme-B (QA16A02, 372226), PE-Fire 700, anti-mouse CD4 (GK1.5, 100484), APC/Cy7 anti-mouse TNFα (MP6-XT22, 506344), PE-Cy7 anti-mouse KLRG1 (2F1/KLRG1, 138416), PerCP-Cy 5.5 anti-mouse CD62L (MEL-14, 104432), BV605 anti-mouse CD127 (A7R34, 135025), Pacific blue anti-mouse CD69 (H1.2F3, 104524), Pacific blue anti-mouse CD45.1 (110722), APC anti-mouse PD1 (RL388, 109111), BV711 anti-mouse PD1 (29F.1A12, 135231), PE anti-mouse IFNγ (XMG1.2, 505808), FITC anti-mouse TNFα (MP6-XT22, 506304), Pacific Blue anti-human CD45 (HI30, 982306), APC/Cy7 anti-human CD8 (RPA-T8, 344713). BUV563 anti-mouse LAG3 (C9B7W, 741350), BUV615 anti-mouse CD69 (H1.2F3, 751593), BUV805 anti-mouse CXCR3 (173, 748700), BV421 anti-mouse EOMES (X4-83, 567166), BV480 anti-mouse CD45.1 (A20, 746666), Alexa-Fluor 488 anti-mouse TCF1 (S33-966, 567018), Alexa-Fluor 647 TOX (NAN448B, 568356), BUV 496 anti-mouse Ly-108 (13G3, 750046), APC/Cy7 anti-mouse CD44 (IM7, 560568) and BUV805 anti-mouse CD8α (53-6.7, 612898) were acquired from BD Biosciences. BUV395 anti-mouse CD44 (IM7, 363-0441-82), PerCP-Cy 5.5 anti-mouse IL-2 (JES6-5H4, 45-7021-82), PerCP-eF710 anti-mouse CD27 (O323, 46-0279-42), and PE-Cy7 anti-mouse Tim-3 (RMT3-23, 12-5870-82, eBioscience) were obtained from Thermo Fisher. APC anti-human/mouse Tox (REA473, 130-118-335) was obtained from Miltenyi Biotech. For surface markers, the cells were stained for 30 min on ice in PBS containing 2% FCS and 0.1% sodium azide (cat. S2002, Sigma-Aldrich). To examine intracellular cytokine production, the isolated cells were stimulated with a cell activation cocktail (cat. 5476, TOCRIS) in the presence of monensin (cat. 554724, BD Biosciences) for 4 hr, and stained for intracellular cytokines and transcription factors using a fixation/permeabilization kit (cat. 00-5523-00, Ebiosciences). Fixable viability dye (Thermo Fisher) was used to exclude dead cells. Samples were acquired on a Cytek Aurora (SpectroFlo) or BD LSRII (FACSDiva) flow cytometer and analysed using FlowJo software.

### *In vivo* tumor analysis

For all tumor experiments, 0.5×10^6^ tumor cells were injected subcutaneously into the right flank of naive mice (aged 6–10 weeks). Met was diluted in HBSS to a concentration of 61 μM per 50μl. Mice were injected peri-tumorally for 5 days with either 50 μl Met or HBSS. Tumours were measured with calipers every 2 days and tumor volumes were calculated using the formula: (width^2^ × length)/2. For adoptive transfer experiments, T cells (1 × 10^6^) were transferred i.v. 7 −9 days after tumor injection. For contralateral tumor model, 0.5 × 10^6^ B16 tumor cells were injected subcutaneously into the right flank and left shoulder of the same mouse. Met or HBSS was injected peri-tumorally to the right flank tumor (abdominal). For check-point blockade experiment, 0.5 × 10^6^ MC38 tumor cells were injected subcutaneously into the right flank of WT mice. The mice were given either anti-PD1 antibody (RMP1-14, Bio X Cell) or rat IgG2a (2A3, Bio X Cell), intraperitoneally twice at a dose of 150 mg/kg in 100 μl PBS at day 9 and 12. Met or HBSS was given peri-tumorally for 5 days from day 9 to 13. For the CAR-T tumor model, 0.5 × 10^6^ F420 tumor cells (derived from singly floxed *p53+/F-Col2.3* transgenic mice^2^); were injected subcutaneously into the right flank of *Rag1*^-/-^ mice. 5 × 10^6^ B7-H3 CAR-T cells (see below) were transferred i.v. 8-10 days post implantation. Tumour size was measured every 2-4 days.

### Generation of mouse CAR-T cells

Generation of mouse B7-H3-CAR constructs and CAR-T generation have been previously described^3^. Briefly, HEK293T cells were transiently transfected with the CAR-encoding plasmid, the Peg-Pam plasmid encoding MoMLV gag-pol, and a plasmid encoding the VSVG envelope. Viral supernatants were collected at 48 h and filtered with a 0.45m filter. The virus was then used to transduce the GPE86 producer cell line. B7-H3-CAR-expressing GPE86 cells were stained with Alexa Fluor-647 anti-human IgG, F(ab′)2 fragment antibody (cat. 109-606-006, Jackson ImmunoResearch) and sorted using the BD FACSAria III system.

Naive CD8^+^T cells from 6–8 weeks old CD45.1^+^ mice were activated with plate-bound anti-CD3ε (1 μg/ml, 145-2C11, cat. BE0001, BioXcell), anti-CD28 (2 μg/ml, 37.51, cat. BE0015, BioXcell) and 50 U ml^−1^ rhIL-2 (cat. 200-02, Peprotech). On day 2 post activation, activated CD8^+^ T cells were transduced with retrovirus expressing B7-H3-CAR on retronectin-coated (cat. T100B, Takara) non-tissue-culture-treated plate in complete RPMI medium supplemented with 50 U/ml rhIL-2. At 48 h after transduction, CAR-T cells were collected and expanded in the presence of 50 U/ml rhIL-2 for another 3 days before adoptive transfer into tumor-bearing mice.

### Tumour-infiltrating lymphocyte isolation

B16 tumors were harvested at the indicated time points and digested using a digestion cocktail (0.1% Collagenase IV and 0.01% DNaseI) for 5 min at 37 °C. Digested tumors were passed through a 70m strainer and resuspended in PBS plus 2% dialyzed FCS and processed for analysis and/or sorting by flow cytometry, as described above.

Patient samples were collected following surgery and were immediately processed. Tumour samples were dissected into small pieces and digested at 37°C with digestion cocktail for 40 min. Digested tumor samples were physically disrupted and passed through 70μ filter. Samples were washed with PBS-2% dialyzed FCS twice and stained for CD45, CD3ε and CD8 and sorted for CD8^+^ T cells by flow cytometry. Purified T cells were washed once with 1% saline and flash frozen in liquid nitrogen. Samples were collected upon informed consent and procedures approved by the IRB of St. Jude Children’s Research Hospital.

### Immunofluorescence staining and imaging

CD8^+^ T cells were immobilized in 8-well poly L-lysine-coated IBIDI chambers and stimulated by the addition of CD3/28 Dynabeads (cat. 11452D, Thermo Fisher) for 30 min, followed by 10 min of fixation with 4% paraformaldehyde (PFA) (cat. 15710, Electron Microscopy Science) at 37°C. The cells were washed once with TBS (50 mM Tris, 100 mM NaCl, pH 8.0) and permeabilized with permeabilization buffer (50mM Tris, 100 mM NaCl, pH 8.0, 0.3% (v/v) Triton X-100) for 20 min at room temperature. Cells were washed again with TBS before blocking with TBS containing 2% BSA (Cat. 001-000-182, Jackson ImmunoResearch) for 60 min at room temperature. The cells were stained overnight at 4 °C with the following primary antibodies: anti-NFAT1 (1:200, cat. 4389, Cell Signaling Technology), anti-NFAT2 (1:200, cat. D15F1, Cell Signaling Technology) and anti-mono and dimethyl arginine (1:500, cat. ab412, Abcam). The samples were washed twice with TBS and incubated for 1 h at room temperature with the following antibodies: anti-rabbit Alexa Fluor plus 595 (1:1,000, cat. A-11012, Thermo Fisher), anti-mouse CD8-APC (1:500; cat. 100712, BioLegend), Alexa Fluor 488 Phalloidin (1:1000, cat. A12379, Thermo Fisher) and Hoechst 33258 (1:1000, cat. H3569, Thermo-Fisher). Samples were then imaged by Marianas spinning-disc confocal microscopy (Intelligent Imaging Innovations), using the Prime95B sCMOS camera, and the 405, 488, 561 and 640 nm laser lines. Sum fluorescence intensities were analyzed using Slide Book 6 (Intelligent Imaging Innovations).

### Immunoblotting

T cells were washed once with ice-cold PBS and immediately lysed with 4x laemmli buffer (Bio-Rad, cat. 1610747). Cell lysates were separated by SDS-PAGE and blotted with anti-Kcnn4 antibody (cat. PA5-33875, Thermo Fisher) and HRP-conjugated anti-Actin (cat. sc4777-HRP, Santa Cruz Biotechnology). Images were developed using Clarity Western ECL substrate (cat. 1705061, Bio-Rad) and acquired on Bio-Rad Chemi-Doc.

### Amino acid measurement

Subcutaneous tumors were isolated and processed as described above. Sorted T cells were washed once with ice-cold saline, flash-frozen with liquid nitrogen, and stored at –80°C before analysis. Cell pellets were extracted using 750 µl of methanol/acetonitrile/water (5:3:2, v/v/v) and the supernatant was dried by lyophilization. Aliquots of 20-50 µl from plasma and TIF were extracted with at least a 15-fold excess volume of the methanol/acetonitrile/water solution, and the supernatant was then collected and dried by lyophilization. Dried extracts containing the hydrophilic metabolites were dissolved in 30 µl of water/acetonitrile (8:2, v/v), and 10 µl was used to derivatize amino acids as described ^4^ with some modifications. The samples were placed in glass autosampler vials, and 35 µl of sodium borate buffer (100 mM, pH 9.0) was added and mixed by pipetting. 10 µl of the 6-aminoquinolyl-N-hydroxysuccinimidyl carbamate (AQC, 10 mM in acetonitrile)-derivatizing reagent (Cayman Chemical) was added, the vial was sealed, mixed by vortexing, and then incubated at 55°C for 15 min. The vial was cooled to room temperature and then 1 µL was analyzed by liquid chromatography with tandem mass spectrometry (LC-MS/MS). An ACQUITY Premier UPLC System (Waters Corp) was used for the LC separations, using a non-linear gradient condition as follows: 0-0.4 min 3% B; 0.4-8 min 3 to 96% B (using Curve #8 of the inlet condition in MassLynx™); 8-12 min 96% B; 12-12.5 min 96 to 3% B; 12.5-14 min 3% B. Mobile phase A was water supplemented with 0.15% acetic acid, and mobile phase B was acetonitrile with 0.15% acetic acid. The column used was an Accucore C30 (50 × 2.1 mm, 2.6 μm) (Thermo Fisher Scientific) operated at 50°C. The flow rate was 300 μl/min and the injection volume used was 1 μl. All LC/MS solvents and reagents were the highest purity available (water, acetonitrile, acetic acid, boric acid, sodium hydroxide) and were purchased from Thermo Fisher Scientific. A Xevo TQ-XS Triple Quadrupole Mass Spectrometry (TQ-XS) (Waters Corp) equipped with a multi-mode ESI/APCI/ESCi ion source was employed as detector. The TQ-XS was operated in positive ion mode using a multiple reaction monitoring mass spec method (MRM).

The MRM conditions were set to a minimum of 15 points per peak with an automatic dwell time. The operating conditions of the source were: Capillary Voltage 3.8 kV, Cone Voltage 40 V, Desolvation Temp 550°C, Desolvation Gas Flow 1000 L/h, Cone Gas Flow 150 L/h, Nebuliser 7.0 Bar, Source Temp 150°C. Authentic amino acid standards were purchased from Sigma-Aldrich and used to establish MRM conditions and calibration curves. The monitored parent/daughter ions, fragmentation collision energy (CE), and retention time window for each amino acid are listed in Supplementary Table 4. The acquired MRM data was processed using the software application Skyline 21.2 (MacCoss Lab Software).

### *In-silico* model building of KCa3.1(SK4)-Calmodulin complex and simulation

To study the impact of SK4s arginine dimethylation on the binding of calmodulin, we simulated the SK4 channel:Calmodulin complex in a membrane layer via all-atom molecular dynamics (MD), which is considered a computational microscope^5^. The Cryo-EM structure of the human SK4 channel:Calmodulin complex state (PDB:6CNN^6^) was used as the starting structure. Thisa tetramer compleconsistsng of four SK4 monomers, four calmodulins (CAMs), three calcium ions bound to each calmodulin (CAM) (total 1,2) and four K^+^ ions bound to the channel selectivity filter.

Initially, protein was prepared was using the molecular operating environment (MOE) software. The missing residues in the cryo-EM structure were modeled based on the human AlphaFold structure (AF-015554-F1). Protonate3D module in MOE was used to assign appropriate protonation states at pH 7.4 and then energy minimized. Further, simulation system was prepared using CHARMM-GUI web server^7–9^. Protein was placed in the POPE membrane bilayer and protein orientation was obtained using the PPM web server^10^. The protein-membrane bilayer complex was immersed in the TIP3P^11^ water box of size ≈150 × 150 × 230 Å^3^ with an edge distance of 40 Å on all sides. KCl salt was added to balance the charges and maintain a physiological salt concentration of 0.15M. Overall, each wildtype simulation system contains four SK4 monomers, four CAMs, 12 Ca^2+^ ions, ≈355 K^+^ ions, ≈367 Cl^-^ ions, ≈618 POPE lipids, and ≈128,144 TIP3P waters. The total numbers of atoms in this system are ≈498,944. Two more systems were generated via a protocol similar to that explained above, except that in one system, R352 of the SK4 monomer was symmetrically dimethylated and in the second system, R352 was asymmetrically dimethylated. Dimethylation was introduced using the CHARMM-GUI web server. Note that arginine’s at position 352 in all four SK4 monomers were dimethylated in both systems.

Further, all three systems were simulated using AMBER20/22 MD software^12^. The MD simulation was performed in three steps. In step-1, energy minimization was carried out to remove potential steric clashes in two stages. In stage 1, each system was energy-minimized for 5000 steps, while positional restraints were applied to the protein and membrane: the first 2500 steps with the steepest-descent algorithm and the remaining 2500 steps using the conjugate gradient algorithm^13^. In stage 2, the second round of minimization was carried out with all restraints removed in 50,000 steps. The steepest-descent algorithm^14^ was used for the first 25,000 steps, while the conjugate gradient algorithm was used for the remaining 25,000 steps. In step-2, equilibration was carried in six stages: (1) equilibrated for 125 ps with a time step of 1fs while applying positional restraints on protein (K=10 kcal/mol.Å^2^) and membrane (K=2.5 kcal/mol.Å^2^), (2) equilibrated for 125 ps with a time step of 1fs w while applying positional restraints on the protein (K=5.0 kcal/mol.Å^2^) and membrane (K=2.5 kcal/mol.Å^2^), (3) equilibrated for 125 ps with a time step of 1fs while applying positional restraints on the protein (K=2.5 kcal/mol.Å^2^) and membrane (K=1.0 kcal/mol.Å^2^), (4) equilibrated for 500 ps with a time step of 2 fs while applying positional restraints on the protein (K=1.0 kcal/mol.Å^2^) and membrane (K=0.5 kcal/mol.Å^2^), (5) equilibrated for 500 ps with a time step of 2 fs while applying positional restraints on the protein (K=0.5 kcal/mol.Å^2^) and membrane (K=0.1 kcal/mol.Å^2^) and (6) equilibrated for 500 ps with a time step of 2 fs while applying weak positional restraints on the protein (K=0.5 kcal/mol.Å^2^) and restraints on the rest of the system were completely removed. The first two equilibration simulations were performed under NVT conditions, whereas the last four equilibration simulations were performed under NPT conditions. In step-3, production simulations were carried out under periodic boundary conditions using the NPT ensemble, during which the temperature (310 K) and pressure (1 atm) were kept constant. Three replicates of each system were simulated, and each replicate was simulated for ≈1 µs. Langevin integrator and Monte Carlo barostat were used^15,16^ to conduct NPT simulations, whereas the SHAKE algorithm was used to restrain bonds with hydrogens^17^. Particle-mesh Ewald (PME) method was used to treat long-ranged electrostatic interactions^18^ and the nonbonded cutoff was 12Å (force switching = 10Å). CHARM36 all-atom additive force field^19,20^ was used to treat the entire system. Overall, nine MD simulations were conducted across three systems, with each system comprising three replicates that each ran for approximately 1µs. Consequently, the total MD simulation data accumulated for analysis amounts to approx. 9 µs, derived from the sum of simulation times for all systems and replicates (3 systems X 3 replicates X ≈1µs/per replicate = ≈9µs). Two frames per nanosecond were used for the data analysis. Visual molecular dynamics (VMD)^21^ was used for visualization and graphic generation, whereas Tcl was used for data analysis. Gnuplot was used to generate the time series plots.

### ATAC-seq

OT-I CD8^+^ T cells were activated with 2.5ng/ml SIINFEKL in either 0.1 mM Met or 0.03 mM Met with restoration of Met to 0.1 mM in 0.03 mM condition post 30 min of activation and incubated for 24 hrs. KCa3.1^WT^ and KCa3.1^R350A^ cells were activated by 2.5ng/ml SIINFEKL for 24 hrs. Cells were processed for ATAC-Seq after 24 hrs of activation. Briefly, 50,000 cells were washed with 1 ml cold PBS by centrifugation at 500 g for 5 min at 4°C. Cells were then resuspended in 50 ml cold cell lysis buffer (10 mM Tris pH 7.4, 10 mM NaCl, 3 mM MgCl_2_, and 0.1% NP-40). Isolated nuclei were pelleted at 1,000 × g for 10 min at 4°C, resuspended in 50 μl transposition reaction mixture (25 μl TD buffer, 2.5 μl TDE1, 22.5 μl ddH_2_O) (20034197, Illumina), and incubated at 42°C for 45 min. DNA was purified using a MiniElute PCR purification kit (cat. 28004, Qiagen). Samples were then processed for library barcoding and amplification with Q5 High-Fidelity 2× Master Mix (cat. M0492S, NEB). Prepared libraries were sent for sequencing after quantification using Qubit and size distribution as determined using an Agilent 4200 TapeStation.

### Analysis of ATAC-seq data

Paired-end sequencing reads were trimmed using Trim Galore (version 0.5.0) (https://github.com/FelixKrueger/TrimGalore) with default parameters. Reads were aligned to the reference mouse mm10 assembly using Bowtie 2 (version 2.3.5.1)^22^ with settings --end-to-end -- very-sensitive -X 2000. The resulting alignments, recorded in a BAM file, were sorted, indexed, and marked for duplicates with Picard MarkDuplicates function (version 2.19.0). Afterward, the BAM file was filtered with SAMtools (version 1.9)^23^, BamTools (version 2.5.1)^24^ and scripts of nf-core/chipseq^25^ to discard reads, mates that were unmapped, PCR/optical duplicates, not primary alignments, mapped to multiple locations, or mapped to ENCODE blacklisted regions^26^; only reads mapped in proper pair were kept (-F 1804 -f 2 -q 30). The alignments were shifted by deepTools alignmentSieve^27^ with --ATACshift and default parameters. Nucleosome-free reads (fragment length < 109 bp) were separated from the BAM files with SAMtools (version 1.9). MACS (version 2.1.2)^28^ was used to call peaks from the BAM files with narrowPeak setting, --extsize 200, and recommended mappable genome size (default value for other parameters). Chromatin accessibility signal was normalized by scaling to 1 million mapped reads using BEDTools (version 2.27.1)^29^ and bedGraphToBigWig (version 377)^30^ and visualized as heatmaps using deepTools plotHeatmap (version 3.2.1)^27^.

Differentially accessibility analysis was performed using DiffBind^31^ (v.3.4.7) (summit = 50, normalized by trimmed mean of M values^32^ (TMM) approach, library size estimated by reads mapped to “background” regions, the genomic regions (binned into 15000 bp) overlapped with peaks). Peaks were annotated to the nearest genes with annotatePeak function in R package ChIPseeker^33^ (v.1.30.3) at the gene level using the default parameters. Genes were ranked by fold change (log2 transformed) of associated peaks located within +/- 1.5 kb of transcription start sites (if a gene was associated with multiple peaks, the mean was used). Gene Set Enrichment Analysis^34^ (GSEA) was then performed against MSigDB collections with R package clusterProfiler^35^ (v.4.2.2). The common differentially accessible regions (DAR) between ATAC-Seq of KCa3.1WT and KCa3.1R350A and T cells activated in 0.1 mM or 0.03 mM Met were obtained by the overlapping of the DARs (at least 1 bp). Enrichment analysis was performed using the over-representation approach against MSigDB using the R package clusterProfiler.

### CUT&RUN

CUT&RUN experiments were performed as previously described^36^_with slight modifications. Purified OT-I CD8^+^ T cells were activated with 2.5ng/ml SIINFEKL in 0.1 mM and 0.03 mM Met for 30 min. Met was restored to 0.1 mM post 30 min and cultured for 24 hrs. Cells were washed with cold PBS and dead cells were removed using Easysep dead cell removal kit (cat. 17899, StemCell). Cells (2 × 10^5^) were washed twice with wash buffer (20 mM HEPES (H3375, Sigma-Aldrich], 150 mM NaCl (cat. AM9760G, Invitrogen), 0.5 mM spermidine (cat. S0266, Sigma-Aldrich), and protease inhibitor cocktail (cat. 5056489001, Sigma-Aldrich)), resuspended and bound to concanavalin-A coated magnetic beads (cat. BP531, Bang Laboratories). The resuspension buffer was removed using a magnetic stand and beads were resuspended in 100 μl antibody buffer (20 mM HEPES, 150 mM NaCl, 0.5 mM spermidine, 0.01% digitonin (cat. 300410, Millipore), 2 mM EDTA (cat. AM9260G, Invitrogen), and protease inhibitor cocktail). Primary antibodies (1:100) were added to the samples and incubated overnight at 4 °C. Next, samples were washed twice with cold Dig-wash buffer (20 mM HEPES, 150 mM NaCl, 0.5 mM spermidine, 0.01% digitonin, and protease inhibitor) and pAG-MNase (cat. 123461, Addgene) was added and rotated at 4 °C for 1 h. Samples were washed twice and resuspended in 50 μl Dig-wash buffer and CaCl_2_ (2 μl of 100 mM, cat. 2115, Sigma-Aldrich) was added followed by brief vortexing and incubation on ice for 30 min. Next, 50 μl of 2× STOP buffer (340 mM NaCl, 20 mM EDTA, 4 mM EGTA (cat. E3889, Sigma-Aldrich) and 100 μg/ml RNase A (cat. EN0531, Thermo Fisher), and 50 μg/ml GlycoBlue (cat. AM9515, Invitrogen)) was added and mixed by gentle vortexing and incubated for 30 min at 37 °C to release CUT&RUN fragments. Fragmented DNA was purified using the NEB Monarch PCR&DNA Cleanup Kit (cat. T1030S, New England Biolabs). DNA libraries were prepared using the NEBNext Ultra II DNA Library Prep Kit (cat. E7645S, New England Biolabs) and purified with AMPure SPRI beads (cat. B23318, Beckman-Coulter). Prepared libraries were quantified using Qubit and size distribution was determined with a Agilent 4200 TapeStation analysis before paired-end sequencing.

### CUT&RUN data processing

Paired-end sequencing reads were trimmed using Trim Galore (v.0.5.0) (https://github.com/FelixKrueger/TrimGalore) with default parameters. The reads were then aligned to the reference mouse mm10 assembly using Bowtie 2 (v.2.3.5.1)^22^ with settings --end-to-end --very-sensitive --no-mixed --no-discordant -q --phred33 -I 10 -X 700. The resulting alignments, recorded in the BAM file, were sorted, indexed, and marked for duplicates using the Picard MarkDuplicates function (v.2.19.0) (Picard toolkit. *Broad Institute GitHub Repository,* 2019). Next, the BAM file was filtered with SAMtools (v.1.9)^23^, BamTools (v.2.5.1)^24^ and scripts of nf-core/chipseq^25^ to discard reads, including mates that were unmapped, PCR/optical duplicates, non-primary alignments, reads that mapped to multiple locations or to ENCODE-blacklisted regions^26^, or with more than 4 mismatches (-F 0×004 -F 0×008 -F 0×0100 -F 0×0400 -f 0×001 -q 1). MACS (v.2.1.2)^28^ was used to call peaks from the BAM file with IgG control and recommended mappable genome size (the default values were used for the other parameters). NarrowPeak mode was used for NFAT1, H3K27me3, and H3K4me3. Binding signal was normalized by scaling to 1 million mapped reads using BEDTools (version 2.27.1)^29^ and bedGraphToBigWig (version 377)^30^ and visualized as heatmaps using deepTools plotHeatmap (version 3.2.1)^27^

In the CUT&RUN experiment with spike-in E. coli, two modifications were made: i) a hybrid reference of mouse mm10 and E. coli ASM584v2, and ii) signals were normalized by scaling to per million reads mapped to E. coli.

Differential binding analysis was performed using DiffBind (v.3.4.7) (summit = 75 (for NFAT1) and 100 (for H3K27me3 and K3K4me3 modifications), normalized by the trimmed mean of M values (TMM) approach library size estimated by reads mapped to “background” regions. Peaks were annotated to the nearest genes with the annotatePeak function in R package ChIPseeker (v.1.30.3) at the gene level using default parameters. Genes were ranked by fold-change (log2 transformed; if a gene was associated with multiple peaks, the mean was used). GSEA was then performed against MSigDB collections with R package clusterProfiler (v.4.2.2).

### RNA-seq

The isolated CD8^+^ TIL were washed once with cold PBS and pelleted at 1500 rpm at 4°C. Total RNA was isolated using the Direct-zol RNA Microprep Kit (cat. R2061, Zymo Research) according to the manufacturer’s instructions and quantified using an Agilent 4200 TapeStation. Libraries were prepared using the KAPA RNA HyperPrep Kit with RiboErase (HMR) (cat. 08098131702, Roche) and purified by AMPure SPRI beads (cat. B23318, Beckman-Coulter). Libraries were quantified and size distribution was determined using a Agilent 4200 TapeStation before paired-end sequencing was performed.

### RNA-seq data processing

Paired-end sequencing reads were mapped by the pipeline of St. Jude Center for Applied Bioinformatics. The reads were trimmed using Trim Galore (version 0.5.0) (https://github.com/FelixKrueger/TrimGalore) with default parameters. Reads were aligned to the reference mouse mm10 assembly plus ERCC spike-in sequences using STAR (version 2.7.5a)^37^. The resulting alignments, recorded in BAM file, were sorted, indexed, and marked for duplicates with Picard MarkDuplicates function (version 2.19.0). Transcript quantification was calculated using RSEM^38^. Differential gene expression analysis was carried out with DESeq2 with Wald test (default parameters)^39^. To examine if global changes in gene expression were present, the RUVg function in the RUVSeq package was used for normalization with ERCC spike-in followed by DESeq2 according to the RUVSeq manual^40^. Gene set enrichment analysis (GSEA) was performed against the MSigDB database^41^ with R package clusterProfiler (version 4.2.2)^35^ with the genes ranked by the Wald statistic from DESeq2 analysis.

### Quantitative Proteomics by tandem mass tag mass spectrometry (TMT-MS)

CD8^+^ T cells were isolated from WT mice and activated with anti-CD3 antibody by crosslinking with IgG for 30 min. Activated and untreated cells were washed once with ice-cold PBS and flash-frozen in liquid nitrogen. Proteomic profiling of the whole proteome and methylome was carried out using a previously optimized protocol^42^ with modifications. Briefly, the cells were lysed in 8M Urea lysis buffer using pulse sonication, and approximately 100 µg of protein per sample was digested with Lys-C (Wako, 1:100 w/w) at 21 °C for 2 h, followed by dilution to reduce urea to 2M, and further digestion with trypsin (Promega, 1:50 w/w) at 21 °C overnight. The protein digests were acidified (trifluoroacetic acid to 1%), centrifuged at 21,000 x g for 10 min at 4°C to remove any insoluble material, desalted with Sep-Pak C18 cartridge (Waters), and dried by Speedvac. Each sample was resuspended in 50 mM HEPES (pH 8.5), TMT labeled, mixed equally, desalted, and fractionated by offline HPLC (Agilent 1220) using basic-pH reverse phase LC (Waters XBridge C18 column, 3.5 μm particle size, 4.6 mm x 25 cm, 180 min gradient, 80 fractions). Fractions were aliquoted (5% v/v) for whole proteome and methylome (95% v/v) profiling. The methylome fractions were further concatenated into 10 fractions for antibody-based sequential mono-methyl-arginine and di-methyl-arginine peptide enrichment.

### Antibody-based Sequential Methylome Enrichment

Each concatenated fraction (approximately 1 mg) was resuspended in 400 µL ice-cold IAP buffer (50mM MOPS, pH 7.2, 10 mM sodium phosphate, and 50 mM NaCl) and centrifuged at 21,000 × g for 10 min at 4°C to remove any insoluble material. The peptides were then incubated with mono-methyl-Arginine antibody beads (mono-me-R, 12235S Cell Signaling Technology) at an antibody-to-peptide ratio of 1:20 (w/w, optimized through a pilot experiment) for 2 hr at 4°C with gentle end-over-end rotation. The antibody beads were then collected by a brief centrifugation, washed 3 times with 1 mL ice-cold IAP buffer and twice with 1 mL ice-cold PBS, while the supernatants were carefully removed and used for the sequential di-methyl-Arginine peptide enrichment (di-me-R symmetric, cat.13563S Cell Signaling Technology). Peptides were eluted from the beads twice at room temperature with 50 µL of 0.15% TFA, dried, and analyzed by LC-MS/MS.

### LC-MS/MS Analysis

Each sample was dried, reconstituted, and analyzed by LC-MS/MS (a CoAnn 75 µm x 30 cm column, packed with 1.9 µm C18 resin from Dr. Maisch GmbH) interfaced with the Orbitrap Fusion MS (Thermo Fisher). LC settings included an 80 min gradient of 15%-40% buffer B (70% ACN, 2.5% DMSO, and 0.1% FA) with buffer A (2.5% DMSO, and 0.1% FA) at a flow rate of ∼0.25 µL /min flow rate. MS settings included data-dependent (3 s cycle) mode with a survey scan in the Orbitrap (60,000 resolution, scan range 410–1600 m/z, 1 × 106 AGC target, and 50 ms maximal ion time), followed by sequential isolation of abundant ions in a 3 s duty cycle, with fragmentation by higher-energy collisional dissociation (HCD, 38 normalized collision energy), and high-resolution detection of MS/MS ions in the Orbitrap (60,000 resolution, 1 × 105 AGC target, 105 ms maximal ion time,1.0 m/z isolation window, and 20 s dynamic exclusion).

### Database Search and TMT Quantification

The MS/MS spectra were searched against the UniProt Mouse protein database (version 2020.04.22) using the COMET algorithm (v2018.013^43^ ) with the JUMP software suite^44^. Search parameters included MS1 mass tolerance of 20 ppm and MS/MS of 0.02 Da, fully tryptic, static mass shift for the TMT16 tags (+304.2071453) and carbamidomethyl modification of 57.02146 on cysteine, dynamic mass shift for Met oxidation (+15.99491), for mono-methyl-Arginine (+14.01565), and for di-methyl-Arginine (+28.0313), maximal missed cleavage (n = 3), and maximal dynamic modifications per peptide (n=5). All matched MS/MS spectra were filtered by mass accuracy and matching scores to reduce false discovery rate (FDR) below 1% for either proteins (whole proteome analysis) or peptides (methylome analysis), based on the target-decoy strategy^45,46^.

TMT quantification analysis was performed as previously reported^47^ with the following modifications: (i) extracting reporter ion intensities from each PSM; (ii) correcting the intensities according to the isotopic distribution of each TMT reagent; (iii) removing PSMs of very low intensities (e.g., minimum value of 1,000 and median value of 5,000); (iv) normalizing sample loading bias with the trimmed median intensity of all PSMs; (v) calculating the mean-centered intensities across samples (e.g., relative intensities between each sample and the mean), (vi) summarizing protein relative intensities by averaging related PSMs; (vii) finally deriving protein absolute intensities by multiplying the relative intensities by the grand-mean of the three most highly abundant PSMs. (vii) Protein fold change and p-values of different comparisons were calculated based on the protein intensities, using a log transformation and moderated t-test with the limma R package^48,49^.

### Quantification and statistical analysis

Data were plotted and analysed using GraphPad Prism (GraphPad Software, v.9.2.0). Statistical significance was calculated using unpaired one- or two-tailed Student’s *t*-tests. Two-way ANOVA was performed to compare tumor growth curves. The log-rank (Mantel–Cox) test was performed to compare the mouse survival curves. For Fig. 7b, a linear mixed effects model with log10(y) as the response variable, group as a fixed-effect predictor variable, and mouse as a random-effect predictor variable was fit to the data. This model represents data with cells as random samples from each mouse. The results showed that the mean log10(y) values of the two mouse groups differed by −0.131 units (95% CI: −0.163, −0.0999; p = 6.98 10^-15^). Statistical significance was set at P < 0.05. Data are presented as mean ± SEM. No statistical methods were used to determine sample size.

